# Sustained growth and rapid dispersal of chitin-associated marine bacteria

**DOI:** 10.1101/2023.06.12.544524

**Authors:** Ghita Guessous, Vadim Patsalo, Rohan Balakrishnan, Tolga Çağlar, James R. Williamson, Terence Hwa

## Abstract

Many biogeochemical functions involve bacteria utilizing solid substrates. However, little is known about the coordination of bacterial growth with the kinetics of attachment to and detachment from such substrates. In this quantitative study of *Vibrio sp. 1A01* growing on chitin particles, we reveal the heterogeneous nature of the exponentially growing culture, comprised of two co-existing subpopulations: a minority replicating on chitin particles and a non-replicating majority which was planktonic. This partition resulted from a high rate of cell detachment from particles. Despite high detachment, sustained exponential growth of cells on particles was enabled by the enrichment of extra-cellular chitinases excreted and left behind by detached cells. The “inheritance” of these chitinases sustains the colonizing subpopulation despite its reduced density. This simple mechanism helps to circumvent a tradeoff between growth and dispersal, allowing particle-associated marine heterotrophs to explore new habitats without compromising their fitness on the habitat they have already colonized.

## INTRODUCTION

While bacterial growth in homogeneous environments has been well characterized^1^, much less is known about growth on solid substrates. Unlike in liquid cultures where nutrients are homogeneously distributed and the population increases exponentially, the temporal characteristics of populations growing in heterogeneous cultures can vary substantially depending on the structure of the environment. This is because temporal and spatial differences in nutrient concentrations may result in the emergence of subpopulations, each adapting to their local environments^2^.

Also, growth on solid substrates complicates our understanding of population fitness, as individuals must not only consume their current substrate to replicate, but they must also successfully seed new habitats to grow further. This tension between growth on the current resource and dispersal from it to find new particles is at the heart of the colonization-dispersal trade-off^3–5^.

While seemingly homogeneous, the ocean offers a highly structured nutrient landscape at the microscopic scale^6–8^. Fecal pellets, organic detritus and dead carcasses (generally termed marine snow)^9, 10^ constitute a large pool of resources that heterotrophic bacteria can utilize as a source of nutrients and energy. One of the major constituents of marine snow is chitin^11, 12^, a biopolymer of GlcNAc molecules that is highly insoluble in water^13, 14^. Chitin is degraded by heterotrophic bacteria such as *Vibrios*^15–18^, which express hydrolytic enzymes called chitinases to break down the long polymeric chains into labile nutrients^19, 20^. This makes chitin a favorable candidate for studying bacterial strategies for utilization of particulate substrates.

Previous studies have characterized isolated components of bacteria-chitin interactions, including enzyme kinetics^19, 21–23^, cell replication^16, 17^, cell attachment^24–26^ and detachment dynamics^27^ as well as motility^28, 29^. However, as we will show, growth on chitin is highly dynamic, involving the simultaneous interaction of these processes. It is thus crucial to integrate these components for the same cells and conditions, in order to build a comprehensive understanding of chitin degradation that bridges cellular descriptions to population-level ones. Such a picture would contribute to informing macroscopic models of carbon cycling^2^ and constraining bacterial contributions therein.

In our study, we characterized the space-dependent growth and dispersal of a natural chitinolytic isolate, *Vibrio sp. 1A01*^26–28^, which is able to utilize chitin particles as its sole source of carbon and nitrogen. Despite the heterogeneous environment, the culture exhibited exponential dynamics, with two subpopulations arising: a replicating minority on the particles and a majority of non-replicating cells, continuously detaching from chitin. We uncovered a population-level scheme involving the use of secreted chitinases, which enabled the particle-associated minority to replicate sufficiently rapidly to drive the exponential increase of both subpopulations. This unexpected behavior is interpreted in light of the marine context in which cells must continuously colonize fresh particles to survive. It provides a novel mechanism through which bacterial populations can circumvent the colonization-dispersal tradeoff.

## RESULTS

### Steady-state growth on chitin with two co-existing subpopulations

*Vibrio splendidus* 1A01 was isolated from a natural community of marine bacteria^30^. When incubated in a minimal medium with chitin flakes as the sole source of carbon and nitrogen, the planktonic component of the culture (Fig. 1a) increased exponentially, at a rate *λ* ≈ 0.06 ± 0.01 ℎ^-1^ (∼12h per doubling; open blue circles Fig. 1b). Upon transferring the planktonic component to a fresh medium with fresh chitin particles, the exponential increase resumed at the same rate (solid circles, Fig. 1b). In comparison, growth was much faster (*r*_max_ = 0.8 ± 0.05 ℎ^-1^, ∼1h per doubling) on GlcNAc, the monomer of chitin (diamonds, Fig. 1b) suggesting additional bottleneck(s) related to chitin utilization.

**Figure 1:**
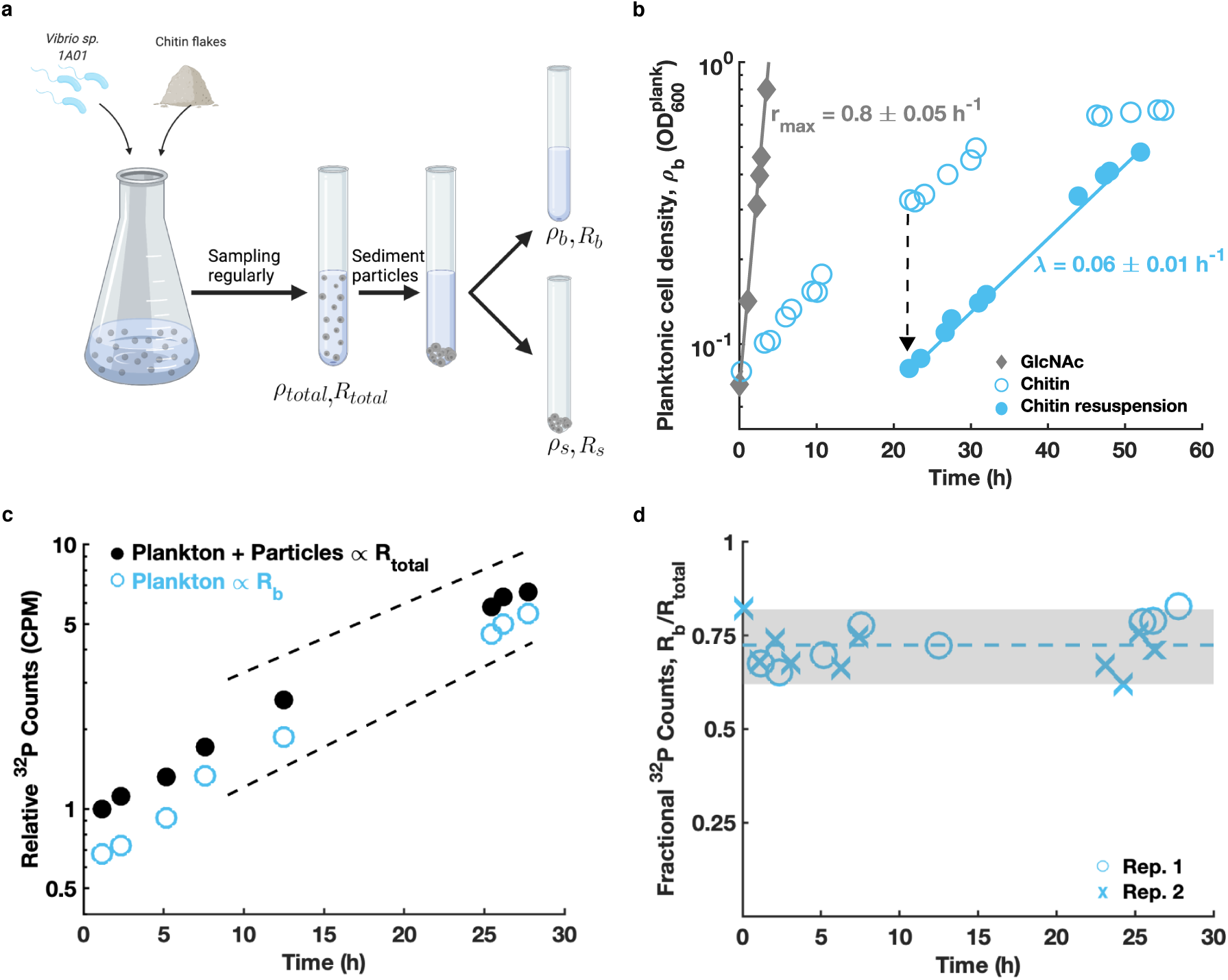
Steady-state growth of two subpopulations on chitin particles. **a)** *Vibrio sp. 1A01* cells were grown with chitin flakes as the sole carbon and nitrogen source. The culture was periodically sampled for measurements. For each sample, the culture was fractionated by sedimenting the chitin flakes and separating them from the planktonic component. Measurements of the planktonic fraction yielded the planktonic cell density (*ρ*_*b*_, Panel b) and planktonic RNA content (*R*_*b*_, Panel c). Measurements of the full culture yielded the total RNA content, (*R*_*total*_, Panel c), and hence the ratio (*R*_*b*_: *R*_*total*_, Panel d). **b)** Open blue circles represent OD600 readings of the planktonic fraction of the chitin culture at various times after inoculation. 24h after inoculation, the planktonic component of this culture was used to inoculate a fresh chitin culture (dashed arrow). Subsequent OD600 measurements are shown as the filled blue circles. The data indicates exponential increase of the planktonic component of the culture at a rate, *λ* = 0.06 ± 0.01 ℎ^-1^ (solid blue line). For comparison, growth of 1A01 on GlcNAc (grey diamonds), exhibits ∼10x faster growth rate (solid grey line). Population increase rates were determined by fitting an exponential model; the error is the standard deviation across six biological replicates. **c)** ^32^P was used to label the cells’ RNA (Methods) in a chitin culture and radioactivity was tracked in samples of the full culture (*R*_*total*_, filled black circles) as well as in planktonic samples (_*b*_, open blue circles). Radioactivity increased exponentially and at approximately the same rate in both. (The upper and lower dashed lines represent population increase rates of *λ* = 0.06 ℎ^-1^and *λ* = 0.07 ℎ^-1^, respectively, and are provided as a guide). **d)** Open blue circles represent the ratio in signal between the two samples in panel c, *R*_*b*_: *R*_*total*_and blue crosses are results from a replicate. The gray area spans the range of our data, between the minimum and maximum estimate. We find *R*_*b*_: *R*_*total*_ ≈ 0.72 ± 0.06 where the error is the standard deviation across all samples. This ratio allows us to deduce the fraction of RNA from particle-associated cells, *R*_*s*_: *R*_*total*_, which is corroborated by direct measurement (Extended Data Fig.1c,d). After adjusting for the cellular RNA amount in each fraction (*R*_*b*_: *ρ*_*b*_ and *R*_*s*_: *ρ*_*s*_) as described in Extended Data Fig. 1b, the ratio of planktonic cells in the culture is deduced: *ρ*_*b*_: *ρ*_*total*_ ≈ 0.75 ± 0.06.

To observe growth of cells on particles, we measured the incorporation of radioactive tracers in biomass: we monitored the incorporation of ^32^P in cells collected from the entire culture, *R*_*total*_(filled circles in Fig. 1c), and from the planktonic component, *R*_*b*_ (open circles). Radioactivity increased at the same exponential rate for both populations (Fig. 1c), similarly to the rate of planktonic OD increase (Fig. 1b). These results establish the coexistence of two exponentially increasing subpopulations: one in the planktonic or **b**ulk state, *ρ*_*b*_, and another on the **s**urface of chitin particles, *ρ*_*s*_. The ratio of radioactive signals in the two samples, *R*_*b*_: *R*_*total*_ (Fig. 1d), together with the RNA contents of the subpopulations *R*_*b*_/*ρ*_*b*_and *R*_*s*_/*ρ*_*s*_(Extended Data Fig. 1a,b), allowed us to estimate the relative abundance of each subpopulation, with *ρ*_*b*_: *ρ*_*total*_ ≈ 0.75 ± 0.06 which was maintained throughout the culture’s growth (Fig. 1d). This is further corroborated by direct measurements of total RNA amounts in each subpopulation (Extended Data Fig. 1c) which yielded an abundance ratio *ρ*_*b*_: *ρ*_*total*_ ≈ 0.8 ± 0.1 (Extended Data Fig. 1d). Thus, while both subpopulations increased exponentially, the majority (75-80%) of the cells were in the planktonic phase. Is the exponential increase in the planktonic fraction due cellular replication or to the detachment of cells from particles?

### Planktonic cells do not replicate despite their increase in density

This increase of the planktonic biomass was surprising since chitin particles are the only source of nutrients in the culture. A possible explanation is that dissolved nutrients such as GlcNAc, generated by surface-associated cells could leak into the medium supporting the replication of planktonic cells. Model calculations taking into account GlcNAc generation on the surface, its uptake and diffusion indicate that this scenario is unlikely beyond a small “screening distance” around the particles for the observed range of cell densities (Supp. Note III-1). Moreover, according to our model this “screening distance”, which depends on the particle density, exponentially decreases, localizing the nutrients to the surface of the particle. Consequently, planktonic cells are highly delocalized and can be treated as effectively being at constant density away from the particles (Supp. Note III-2). Below, we describe a series of experiments establishing the lack of replication of planktonic cells.

First, analyzing the composition of the supernatant by HPLC shows that the concentrations of expected nutrients such as GlcNAc, short GlcNAc oligomers, and acetate are all below our detection limit (Extended Data Fig. 2a). Although low concentrations of dissolved nutrients do not necessarily indicate the lack of cellular replication due to possibly high affinities of *Vibrio* for these substrates, we also found that the supernatant alone does not support the growth of planktonic cells (open diamonds in Fig. 2b).

**Figure 2:**
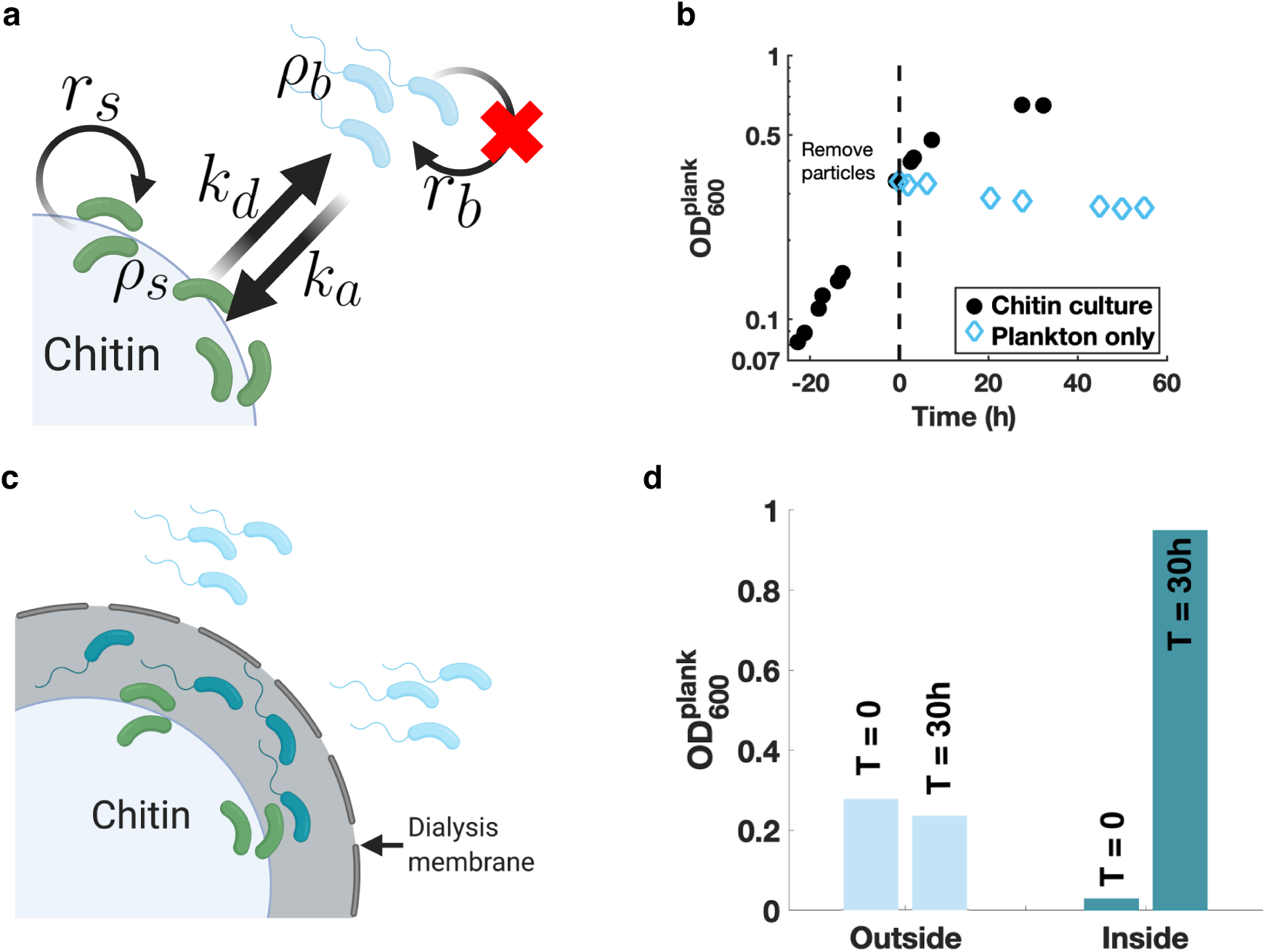
Planktonic cells are not replicating. **a)** A simple population model for the chitin culture: Planktonic and particle-associated cells, with cell densities *ρ*_*b*_ and *ρ*, respectively, replicate with respective rates *r*_*b*_and *r*_*s*_, and exchange with an attachment rate *k*_*a*_ and a detachment rate *k*_*d*_. Both subpopulations increase exponentially and at the same steady-state rate *λ*. The experiments in Panels b and d along with that in Extended Data Fig. 3 lead to the estimate *r*_*b*_ ≈ 0. **b)** Chitin particles were removed from an exponentially increasing chitin culture (black circles) at time 0, and OD600 of the remaining planktonic fraction was tracked (blue diamonds). The OD of the planktonic fraction did not increase in 60 hours, indicating the growth of planktonic cells cannot be sustained without the chitin particles. **c)** From an exponentially increasing chitin culture, chitin particles, colonized with cells (green cells) were placed inside a dialysis bag (grey outline) with a 14kDa molecular weight cutoff, while planktonic cells (blue cells) were left outside of the bag. The pore size allows for small molecules such as GlcNAc to permeate but not enzymes, cells nor chitin particles. Newly released cells from the surface of particles inside of the bag are represented by the teal cells. The OD inside and outside of the bag was monitored after 30 hours to assess the growth of both populations. **d)** Outside the dialysis bag, after 30 hours, the OD of the original planktonic fraction (blue cells in Panel B) did not increase, while inside the bag, the planktonic OD (teal cells) increased ∼3 fold, correspondingly to our expectation for a 30-h period with a population increase rate *λ* ≈ 0.06 ℎ^-1^.

The absence of growth on the supernatant only may be due to the lack of a low but steady supply of dissolved nutrients resulting from chitinolytic activity on the surface. To replicate the situation in our culture, we physically separated planktonic cells from the particles using a dialysis bag (Fig. 2c). While the dialysis bag prevents particles and cells from exchanging, it allows small molecules such as GlcNAc to permeate its membrane (Extended Data Fig. 2b). Tracking the OD of the planktonic component outside of the dialysis bag shows that it did not increase (Fig. 2d), indicating a lack of cellular replication.

Finally, we directly characterized the metabolic activity of the planktonic and particle-associated subpopulations by quantifying their respective RNA synthesis rates. We found that the rate of incorporation of ^3^H-Uridine was 12 times faster in the presence of particles (Extended Data Fig. 3), even though they contained only ¼ of the biomass (Fig. 1d). This suggests that the replication rate of planktonic cells is ∼1/50 of that of cells on particles. Taken together, these experiments indicate that the exponential increase in the density of planktonic cells is not a result of cellular replication.

**Figure 3:**
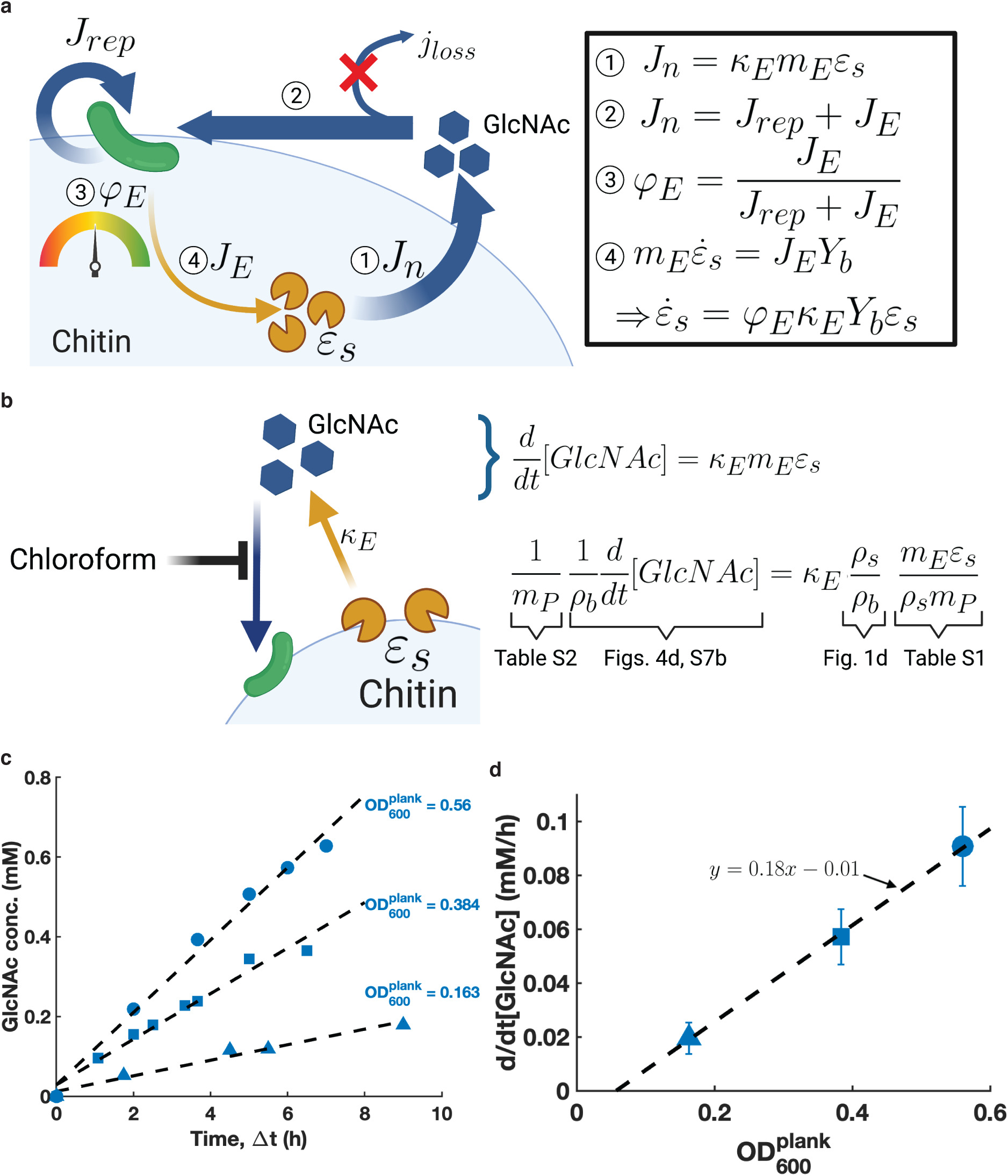
Chitinase activity and the resulting carbon flow. **a)** Model describing chitinase synthesis and labile nutrient generation by chitinases. ① Enzymes attached to the surface of the particles, of concentration *ε*_*s*_ (yellow pacmans), produce GlcNAc molecules (blue hexagons) at a flux *m*_*E*_*κ*_*E*_*ε*_*s*_, where *κ*_*E*_ is the catalytic rate of the chitinases per enzyme mass and *m*_*E*_ the enzyme mass. ② The total flux of generated nutrients *J*_*n*_ is partitioned between cellular biomass production *J*_*rep*_ and enzyme production *J*_*E*_ since the loss of nutrients due to diffusion *j*_*loss*_ is negligible during the exponential phase; see text. ③ The fraction of the total flux *J*_*n*_ allocated towards chitinase synthesis is *φ*_*E*_; this is a key control parameter set by the cells. ④ The rate of enzyme production *m*_*E*_*ε*_*s*_ is given by the flux dedicated to enzyme synthesis *J*, with a nutrient-to-biomass conversion factor, *Y*_*b*_ . Taken together, relations ① through ④ lead to the equation describing the rate of chitinase synthesis 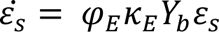, which results in the central equation of our study, Eq.(6), for an exponentially increasing culture limited by labile nutrient generation from chitin. **b)** A simple model of enzyme kinetics. Using a low amount of chloroform (see enzymatic activity in Methods and Extended Data Fig. 6d), we disabled the cells’ ability to uptake labile nutrients. This enabled us to measure the activity of the enzymes (yellow pacmans) by tracking the accumulation of GlcNAc (blue hexagons) concentration in the medium at various time intervals. The specific catalytic rate *κ*_*E*_ can be determined from the rate of accumulation of GlcNAc in the medium (Panels c and d) and the measured chitinase abundance (ED Table 1) through the equations displayed. **c)** GlcNAc concentration is seen to accumulate linearly in time with a rate that varies with the planktonic OD at sampling, reflecting the different amounts of chitinases. The dashed lines represent linear fits to the data, with the flux of GlcNAc generation obtained as the slope of these fits. We note that no GlcNAc oligomers were detected using our procedure; hence we focus on GlcNAc as the primary labile nutrient in this study. We note that the concentrations measured cannot be spatially resolved and are an average in the entire culture. **d)** Flux of GlcNAc generation for samples taken at various planktonic OD shown in Panel c. The error bars represent the 95% confidence intervals of the linear fits done in Panel c. The flux increases linearly with the planktonic OD at sampling. The slope of the linear fit adjusted to the ∼3.3 fold inhibitory effect of chloroform on chitinases (Extended Data Fig. 6d) gives the estimate of 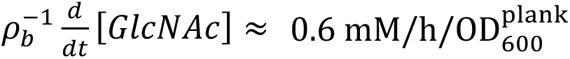 Together with the abundance of chitinases on particles, *m*_*E*_*ε*_*s*_⁄(*ρ*_*s*_*m*_*p*_) ≈ 20% and *m*_*p*_ ≈ 300 *μ*g/ml/OD_600_ (ED Table 1), we find *κ*_*E*_ ≈ 24 ± 5 nmol GlcNAc/µg chitinase/h.

### Rapid shedding of cells from chitin particles supplies the planktonic subpopulation

We hypothesize that, instead, this increase is due to the shedding of cells from particles. Inspecting the culture under a confocal microscope, we found that while chitin particles were only loosely occupied by cells and their surface area was not limiting (Extended Data Fig. 4), cells rapidly attached to or detached from the particle surfaces (Supp. Video), suggesting a dynamic equilibration between the subpopulations of planktonic and surface-associated cells^27^. The detachment dynamics were directly quantified by re-incubating pre-colonized particles in fresh media, and observing the subsequent accumulation of planktonic cells. We found a rapid detachment rate with *k*_*d*_ ≈ 0.18 ℎ^-1^ (Extended Data Fig. 5b).

**Figure 4:**
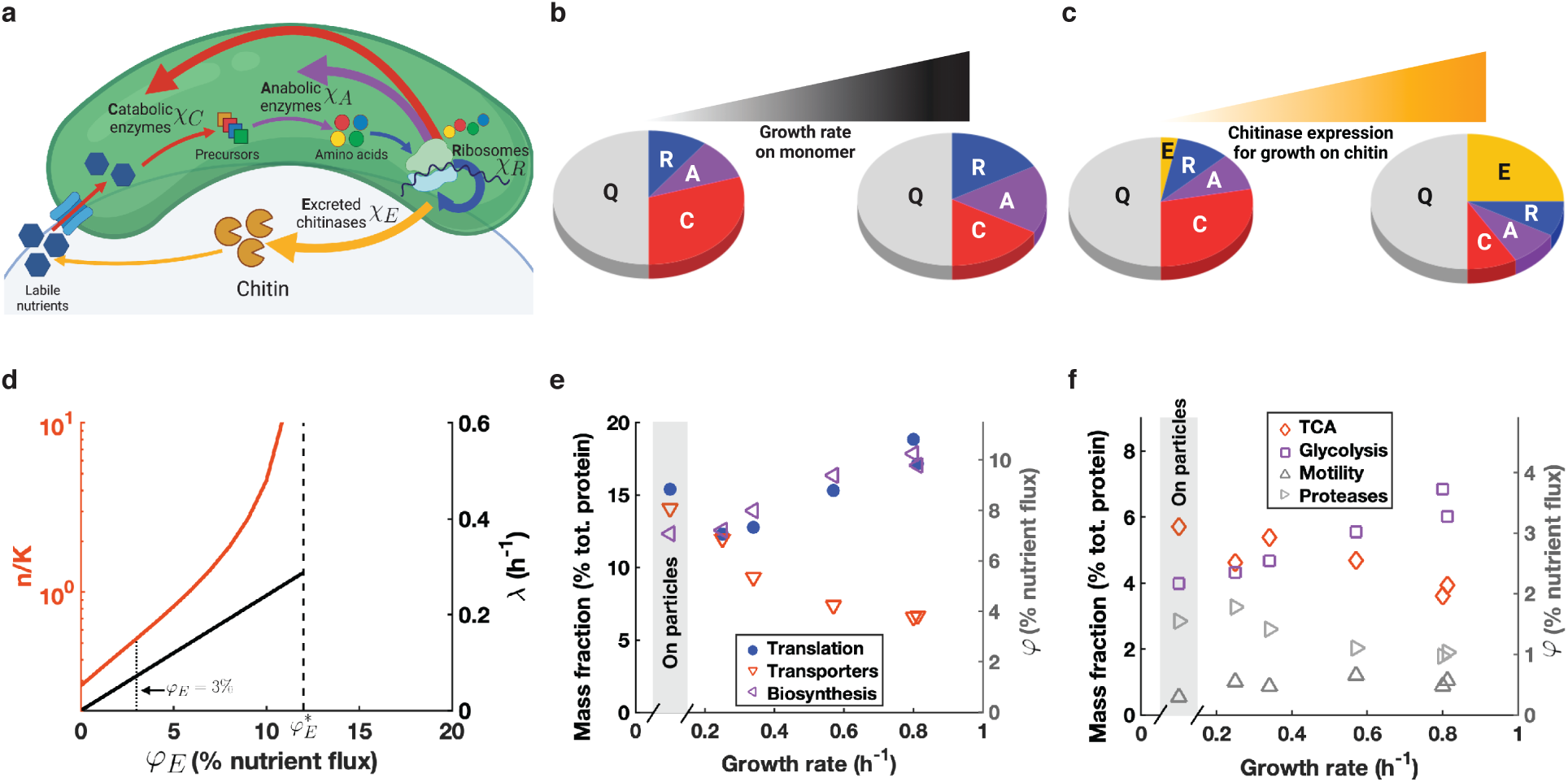
Chitinase expression and proteome allocation. **a)** Illustration of nutrient flow (thin arrows) and protein allocation (thick arrows). Nutrients are generated by **E**xcreted chitinases, taken up and broken down into precursors by **C**atabolic proteins. They are turned into amino acids and nucleotides by **A**nabolic enzymes. Amino acids are assembled by **R**ibosomes and other components of the translational apparatus (R-proteins). C-, A-, and R-proteins are synthesized with fractions *χ*_*C*_, *χ*_*A*_, and *χ*_*R*_of the total protein synthesis flux respectively while chitinases (E-sector) are synthesized with a fraction *χ*_*E*_. The protein synthesis fraction *χ*_i_is related to the nutrient allocation flux *φ*_i_ discussed in the text through the protein:biomass ratio, *b* = *m*_*p*_/*m*_*cell*_(Table S1), by the relation *φ*_i_ = *bχ*_i_⁄[1 − (1 − *b*)*χ*_i_]; see Supp. Note I-1. **b)** Proteome change for growth on good (fast growth) and poor (slow growth) carbon sources based on previous studies of *E. coli* ^38, 39^. Each wedge of the pie-chart indicates the size of the corresponding fraction *χ* introduced in Panel a. On poor carbon sources, protein synthesis is allocated preferentially towards C-proteins (red), reducing the allocation to R- (blue) and A-(purple) proteins. **c)** Even though the synthesis of chitinases (gold wedge), increases the production of nutrients, it reduces the allocation towards the C-, A-, and R-proteins, which adversely affects growth as described by Eq. (7). **d)** Solution to Eqs. 5-7 for various levels of chitinase expression, *φ*_*E*_. The black line indicates the overall population increase rate *λ* (right axis) and the orange line indicates the nutrient concentration on the surface of chitin (left axis). The nutrient concentration diverges for 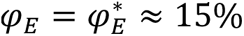 (dashed line). The dotted line represents the measured chitinase fraction of *φ*_*E*_ ≈ 3% (see Extended Data Table 1), which is well below 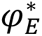. The corresponding nutrient concentration is *n*⁄*K* ≈ 0.4. **e) and f)** Abundances of proteins belonging to various functional groups (as obtained from mass spectrometry) plotted against the growth rate on various carbon sources (Extended Data Fig. 1a, Table S2). We note that the translational group reflects the behavior of the R-sector, the functional groups for biosynthesis and glycolysis reflect the A-sector, while transporters, TCA, and motility reflect the C-sector, analogous to what was found for *E. coli* ^38, 39^. Data in the grey box correspond to samples associated with particles, with chitinases excluded since they are secreted.

**Figure 5:**
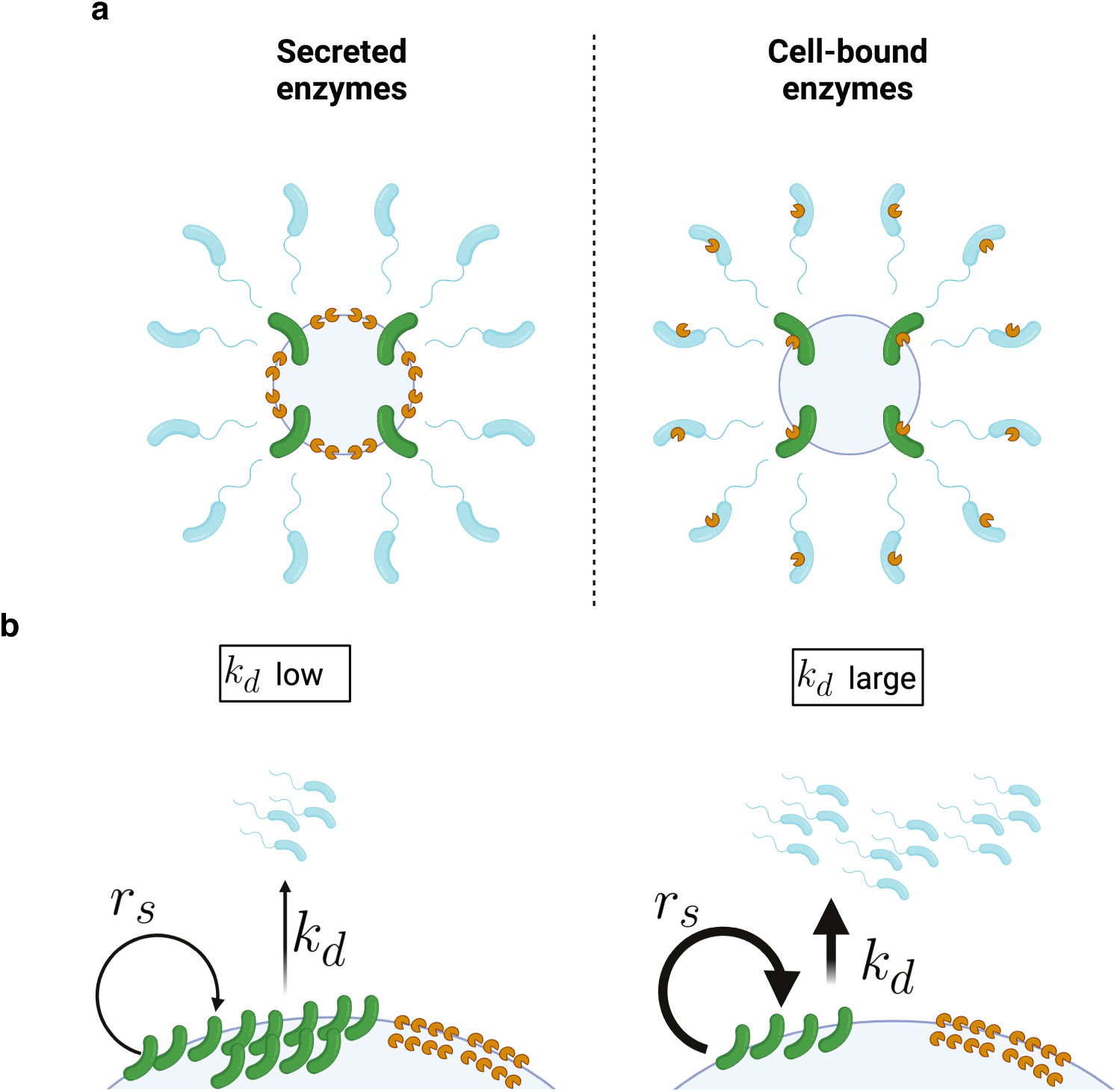
Enzyme secretion is coupled to high dispersal. **a)** Contrasting the cases of secreted and cell-bound enzymes. Secreting enzymes ensures their continued usefulness since they remain attached to the particles even after cells have departed increasing the chitinase-to-cell ratio on the surface of particles. On the other hand, cell-bound enzymes become “useless” as soon as cells depart from the particles resulting in a misallocation of the proteome with a fixed chitinase-to-cell ratio. The difference between these two strategies is modelled in Supp. Note II and Fig. N-II therein. **b)** Enzyme inheritance illustrates the benefits of chitinase secretion. A higher detachment rate results in fewer cells on the surface of the particles, thus increasing the replication rate *r*_*s*_ through the chitinase-to-cell ratio *ε*_*s*_/*ρ*_*s*_. We can see this more formally. In the case of secreted enzymes *χ*_*E*_ = *ε*_*s*_/*ρ*_*total*_. Since the replication rate is proportional to the chitinase amount per cell, this leads to *r*_*s*_ α *χ*_*E*_/(*ρ*_*s*_: *ρ*_*total*_) where *ρ*_*s*_: *ρ*_*total*_ = *λ*/(*λ* + *k*_*d*_) is the fraction of the total population remaining on the particles. For higher detachment rates, *k*_*d*_, this fraction decreases resulting in more chitinases per cell, which translates to a higher replication rate of these cells. This simple feedback between the detachment rate and the replication rate allows for the chitinase expression per cell to be reduced (since the chitinase production is shared by the entire population).

The rate of the reverse process, i.e., the attachment of planktonic cells to chitin particles, is difficult to measure directly due to the difficulty of quantifying a low number of cells on particles. Instead, we estimated this rate using a population model describing the dynamics of two subpopulations of densities *ρ*_*b*_and *ρ*_*s*_, exchanging with attachment and detachment rates, *k*_*a*_ and *k*_*d*_, respectively. Importantly, only particle-associated cells replicate (Fig. 2a), with a rate *r*_*s*_ as shown in the following equations:

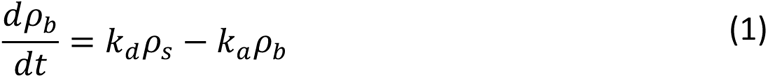

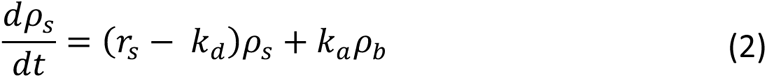

This model admits a dynamic steady state where both sub-populations increase exponentially at the same rate *λ*, with the fraction of the planktonic subpopulation given by *ρ*_*b*_: *ρ*_*total*_ = *k*_*d*_⁄(*λ* + *k*_*a*_ + *k*_*d*_) . Given our measurements of *k*_*d*_ and *λ* above, the planktonic fraction has an upper bound set by *k*_*a*_ → 0, i.e., *ρ*_*b*_: *ρ*_*total*_ ≤ *k*_*d*_⁄(*λ* + *k*_*d*_) ≈0.7 ± 0.1 . Comparison to direct measurements of the planktonic fraction *ρ*_*b*_: *ρ*_*total*_ ≈ 0.75 − 0.8 (Fig. 1d, Extended Data Fig. 1d) suggests that the system is close to this upper bound and thus *k*_*a*_ is negligible.

We validated this by an independent experiment, which established a more stringent bound. By continuously removing the planktonic fraction (shed from the particles) from the chitin culture, we effectively experimentally set *k*_*a*_ ≈ 0 (Extended Data Fig. 5c). We found that the population growth rate obtained in this experiment corresponded to the one measured in the full chitin culture (Extended Data Fig. 5d,e), corroborating the slowness of attachment with a more stringent bound *k*_*a*_ ≪ *λ ρ*_*s*_/*ρ*_*b*_.

### Secreted chitinases are enriched on chitin particles

With attachment kinetics being negligible, the steady state in the chitin culture (Fig. 1c) resulting in exponential growth is obtained by the balance between the replication rate of particle-associated cells, *r*_*s*_, and their detachment rate, *k*_*d*_, with:

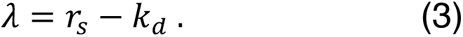

This is the simple statement that cells on particles must replicate fast enough to sustain their rapid detachment rate. Our measured values for *λ* (Fig. 1b,c) and *k*_*d*_ (Extended Data Fig. 5b) lead to the estimate *r*_*s*_ ≈ 0.24 ± 0.04 ℎ^-1^, which although much faster than *λ*, is only one third of the observed maximum growth rate on GlcNAc monomers, *r*_max_ ≈ 0. 8 ℎ^-1^ (Fig. 1b). Thus, the slow rate of population increase (compared to *r*_max_) in our chitin culture resulted from a high detachment rate *k*_*d*_ on top of a moderate replication rate *r*_*s*_.

To understand the source of the reduction of the replication rate from *r*_max_, we turn our attention to the generation of labile substrates such as GlcNAc monomers and oligomers, since their concentrations determine how fast cells replicate. 1A01’s genome is annotated with three chitinases^31^. As different marine bacteria either secrete^16, 17, 32, 33^ their chitinases extra-cellularly or keep them bound to their cell surface^15, 16, 32, 34^, we first established that 1A01 secreted its chitinases (Extended Data Fig. 6a) and that these chitinases were strongly associated with chitin particles (Extended Data Fig. 6b). We next characterized the stability of the secreted chitinases, by isolating them from the cultures’ supernatant (see Fractionation in Methods) and incubating them with fresh chitin particles. There was not a significant difference between the chitinolytic activity measured immediately after the enzymes were collected and the activity 24h later (Extended Data Fig. 6c), thus ruling out significant protein degradation.

### A simple relation connects culture growth to chitinase synthesis

To gain a mechanistic understanding of processes fueling the replication of particle-associated cells, we follow the flow of labile nutrients (collectively referred to as GlcNAc) in the culture. This nutrient flux generated by the secreted chitinases on the particles, *J*_*n*_, is taken up by surface-associated cells for their own replication with a flux *J*_*rep*_ as well as for the synthesis of chitinases with a flux *J*_*E*_(Fig. 3a). In this picture, *J*_*n*_ = *κ*_*E*_*m*_*E*_*ε*_*s*_ with *ε*_*s*_ denoting the concentration of chitinases, *κ*_*E*_ the catalytic rate per enzyme mass and *m*_*E*_ the enzyme mass. In our steady-state culture, the nutrient generation flux as well as the biomass (both cellular and extracellular) increase exponentially, outpacing any loss that may be due to diffusion, *j*_*lossss*_ or transient mismatches in the two fluxes due to heterogeneity in spatial localization of cells and chitinases (Supp. Note III). This is consistent with the observed lack of GlcNAc in the medium as well as the lack of replication by planktonic cells (Fig. 2), leading to *J*_*n*_ = *J*_*rep*_ + *J*_*E*_.

It is convenient to introduce the quantity

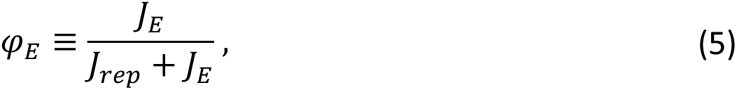

which describes the fraction of the total nutrient flux taken up (*J*_*rep*_ + *J*_*E*_) that is allocated to chitinase synthesis. Since *J*_*E*_ is the flux of nutrients directed towards chitinase synthesis, we have 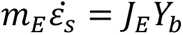 where *Y*_*b*_ is the biomass yield (Table S1). Rewriting Eq. (5) as 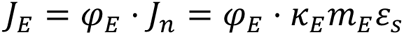, we obtain 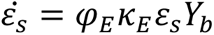, with the steady-state solution:

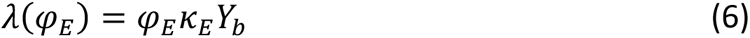

The simple linear relation Eq. (6) makes explicit a key metabolic constraint governing chitin-dependent growth: if the nutrient generated by the chitinases is growth-limiting, then the growth rate of the system is controlled by the allocation towards chitinase synthesis (see Supp. Note I-2). This is analogous to the well-known bacterial growth law describing a linear relation between ribosome content and growth rate, where allocation for ribosome biogenesis is the growth-limiting constraint^35–37^.

To test the validity of Eq. (6), we separately quantified *φ*_*E*_ and *κ*_*E*_ to check if the expression levels and enzymatic properties of the chitinases account for the observed population increase rate. The flux fraction *φ*_*E*_ can be obtained using mass spectrometry. From the mass fraction of chitinases among the total biomass on the particles (∼20%, Extended Data Table 1), we obtain *φ*_*E*_ ≈ 3% (see footnote c, Table S1).

We next determined the *in-situ* catalytic rate of the chitinases *κ*_*E*_. By inhibiting GlcNAc uptake by the cells (Fig. 3b), we tracked the accumulation of GlcNAc in the media at different planktonic ODs (Fig. 3c). The GlcNAc synthesis rate obtained (Fig. 3d), together with the afore-mentioned abundance of chitinases on particles, led us to a specific chitinolytic rate, *κ*_*E*_ ≈ 24 ± 5 nmol GlcNAc/µg chitinase/h (Table S1). Putting these results together, we find the product *φ*_*E*_*κ*_*E*_*Y*_*b*_ to be 0.07 ± 0.01 ℎ^-1^, which is comparable to the directly measured growth rate, *λ* = 0.06 ± 0.01 ℎ^-1^, thus quantitatively supporting Eq. (6).

### Cost and benefit of chitinase expression

Given the maximal replication rate of *r*_max_ ≈ 0.8 ℎ^-1^ on GlcNAc monomers (Fig. 1b) and the measured detachment rate of *k*_*d*_ ≈ 0.2ℎ^-1^ (Extended Data Fig. 5b), the theoretical maximum growth rate of the chitin culture is *λ*_max_ = *r*_max_ − *k*_*d*_ ≈ 0.6 ℎ^-1^ according to Eq. (3). We next examine why the observed growth rate *λ* is 10 times below its upper bound *λ*_max_.

According to Eq. (6), the slowness of the growth rate *λ* may arise from either a slow catalytic rate *κ*_*E*_ or a small allocation towards chitinase synthesis, *φ*_*E*_. The catalytic rate *κ*_*E*_ obtained, which works out to be ∼0.8 GlcNAc molecules/enzyme/s, is actually well within the activity range measured for other metabolic enzymes^23^. This suggests that chitinase expression, (*φ*_*E*_) may be the primary growth-limiting factor.

But how large can the allocation towards chitinase synthesis be? While chitinases are essential for generating labile nutrients (GlcNAc) Eq.(6), their biosynthesis constitutes a burden for processes downstream of GlcNAc generation, including GlcNAc uptake, catabolism, and other biosynthetic processes (Fig. 4a-c). This view^36^ leads us to model these two competing effects of chitinase synthesis on the replication rate *r*_*s*_ of particle-associated cells as :

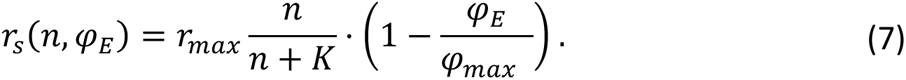

Here *n* is the GlcNAc concentration, *r*_max_ is the aforementioned saturated growth rate on GlcNAc, *K* the Monod constant, and *φ*_max_ ≈ 30 − 40% the maximum fraction of nutrient influx (footnote e, Table S1) dedicated to growth in the fastest growth condition (Supp. Note I).

Eqs. (3), (6), and (7) can be used to solve for the growth rate *λ* for arbitrary values of the allocation *φ*_*E*_. The result (black line in Fig. 4d), shows a positive dependence between the two quantities, despite the cost factor in Eq (7). This is because the benefit of increasing *φ*, as shown by an increasing nutrient concentration *n* (red line in Fig. 4d) strongly compensates for the cost of chitinase synthesis, until *φ*_*E*_ reaches a maximum of 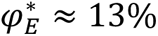 (dashed line in Fig. 4d) when the GlcNAc concentration, *n*, diverges. This corresponds to the replication rate reaching its maximal value: 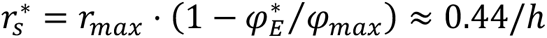 with 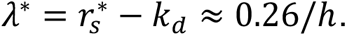. If *φ*_*E*_ is increased beyond 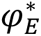, then the replication rate and hence *λ* would decrease according to Eq. (7) since *n* remains divergent. Thus, 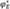 is the “optimal” level of chitinase expression where the growth rate is at its maximum, *λ*^∗^.

### Growth of chitin culture is nutrient-limited due to low chitinase expression

The proteomic tradeoff between chitinases and growth enzymes would result in the reduction of the culture’s maximum growth rate from *λ*_max_ ≈ 0.6/ℎ to *λ*^∗^ ≈ 0.26/ℎ . Nonetheless, *λ*^∗^ is still 3.5 times larger than the observed growth rate *λ* ≈ 0.06/ℎ (Figs. 1b-c). Given the observed chitinase allocation fraction *φ*_*E*_ ≈ 3% is well below the “optimal” allocation 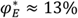, we attribute the reduction in growth rate to a low nutrient concentration as per Eq. (7). For the measured *φ*_*E*_ = 3%, our model predicts a nutrient concentration of *n*⁄*K* ≈ 0.4 suggesting that the slow growth is a consequence of low nutrient concentration, which itself results from a low chitinase expression.

The predicted range of *n*/*K* puts the GlcNAc concentration on the surface of the particles in the *μM* range given the estimated value of *K* (Fig. 4d). Because nutrient limitation is directly reflected in the proteome composition of exponentially growing cells^38, 39^ (Fig. 4b-c), we compared the composition of the proteome of surface-associated cells (excluding chitinases) to that of cells grown under carbon sources that span a range of growth rates from 0.25/h to 0.8/h. We found that a number of proteins whose abundances increased many folds in slow growth had similarly high abundances on particles (Extended Data Fig. 7a). This is also the case for several major groups of metabolic enzymes showing strong growth-rate dependences, with abundances on the particles more closely resembling those on poor carbon sources (open symbols in Figs. 4e, 4f). Similar conclusions are drawn from a more global approach, where the overall proteome of surface-associated cells was more correlated with growth on poor carbon sources (Extended Data Fig. 7c,e). Together, these elements corroborate the finding that the replication rate of particle-associated cells was carbon-limited.

## DISCUSSION

In this study, we characterized the properties of *Vibrio sp. 1A01* utilizing chitin particles. Despite the heterogeneous nature of the substrate, the culture exhibited simple exponential growth. Quantitative experiments established that in this steady-state, the culture segregated into two subpopulations in dynamic equilibrium with each other. Particle-associated cells, although constituting only ¼ of the population, fueled the increase of the entire culture by secreting chitinases, replicating, and eventually shedding from particles, thereby converting to the planktonic sub-population which did not replicate or synthesize chitinases. The overall rate of increase of the culture (*λ* ≈ 0.06 ℎ^-1^) was about 1/3 of the rate of detachment of cells from particles (*k*_*d*_ ≈ 0.2 ℎ^-1^), with a much smaller rate of attachment. Additionally, the replication rate of surface-associated cells, *r*_*s*_ = *λ* + *k*_*d*_ ≈ 0.26 ℎ^-1^, was slow relative to their replication on a saturating amount of GlcNAc (*r*_max_ ≈ 0.8 ℎ^-1^). These results are surprising with respect to the canonical scenario for chitin degradation: instead of attaching and consuming the particles until they’re nearly exhausted, 1A01 preferentially detached from them, even though they are the sole source of nutrients in the culture. We expect this preferential detachment of 1A01 to hold in the ocean as well, where the particle density is much lower^14^ making attachment even less favored.

Remarkably, the small fraction of particle-associated cells was able to drive the exponential increase of the entire population. We analyzed this phenomenon in light of a mechanistic model of chitin utilization, which led to a central relation (Eq.(6)) linking chitinase abundance and activity to growth rate. Given the specific chitinase activity measured (*k*_*ε*_∼1 GlcNAc/enzyme/s which was not particularly low^23^), we expect that increasing chitinase production would result in an increase of growth rate until an upper bound *λ*^∗^ = 0.26 ℎ^-1^ (Fig. 4d). We attribute the slow growth observed, *λ* ≈ *λ*^∗^/4 to the low level of chitinase excretion, estimated to be *φ*_*E*_ ≈ 3% of the total nutrient flux. This low level of chitinase production could be a result of a proteomic cost not probed by this analysis (e.g, secretion machinery), or a strategy to mitigate “cheating” (see below) since higher chitinase expression would result in higher nutrient concentration (Fig. 4d), making it more accessible to other species. Alternatively, slow growth may be favored for ecological reasons in the context of sinking POM (see Extended Data Fig. 8).

The extra-cellular secretion and release of precious, growth-limiting resources is commonly thought of as wasteful from the perspective of the secreting individual as it results in the production of public goods that may escape capture and promote “cheating”. As a way of bypassing this problem, organisms may “privatize” their resources by binding them to the cell surface^16, 40–42^ such that nutrients generated are immediately taken up. Our analysis in Supp. I-2 suggests that such a strategy would be inefficient from a resource allocation perspective as it would require chitinase expression to increase 4-fold, to *φ*_*E*_ ≈ 12%, to support the same growth rate, *λ* (Supp. Note I-4). This inefficiency arises because cells with bound enzymes would carry their enzymes with them as they detach from particles, after which these enzymes become “useless” (Fig. 5a). This highlights a fundamental conflict between “privatizing” enzymes to prevent “cheating” and secreting enzymes to enable rapid dispersal while maintaining growth.

Conversely, the coupling between enzyme secretion and rapid dispersal is advantageous (Fig. 5b). This is because for the same level of chitinase expression, increase in cell detachment leads to fewer cells on particles, thus higher enzyme-to-cell ratio, and higher flux of nutrient generation and higher rate of cell replication on particles. This non-intuitive effect, that increasing dispersal leads to higher on-particle replication rate, is a result of cells on particles “inheriting” enzymes synthesized by their departing relatives. This mechanism underlies the key relation, Eq. (6), which dictates that the population increase rate is independent of the cell detachment rate (up to the maximal possible rate *λ*^∗^). This enzyme inheritance mechanism ensures that the remaining on-particle colonies maintain high replication rates despite large amount of dispersal. By decoupling cellular replication from nutrient production through the extra-cellular release of enzymes, the population, as a whole, is able to circumvent the colonization-dispersal tradeoff that individual cells are subjected to.

Through quantitative characterization of bacterial growth and protein allocation on and away from particles for *Vibrio sp. 1A01*, this study has revealed a simple molecular trick, the release and inheritance of chitinases, which decouples the commonly held trade-off between growth and dispersal at the population level. These two population traits can instead be independently set by various environmental factors (Extended Data Fig. 9), illustrating the intricate interplay between bacterial strategies of substrate utilization and population dynamics in spatially structured nutrient landscapes.

## METHODS

### Strain

*Vibrio sp.* 1A01 was previously isolated in Datta et. al (2016)^30^ from an enrichment of coastal seawater (Nahant, MA) on chitin beads and obtained from the Cordero lab. It was maintained in glycerol stocks. Colonies were streaked on Marine Broth 2216 with 1.5% agar plates.

### Growth experiments

Experiments were carried out in three steps: seed culture in Marine Broth 2216, pre-culture and experimental culture in the identical minimal medium. For the seed culture, single colonies were inoculated into liquid Marine Broth 2216 media until they reached a sizeable density and then transferred to the pre-culture media with either chitin flakes or the designated substrate as a carbon source. Cells were incubated in the pre-culture medium for about 10 doublings (4 days for chitin cultures) and then transferred to the experimental culture for all assays. For chitin cultures, particles were sedimented (see methods below), and the planktonic fraction only was used to begin a new experimental culture.

### Culture conditions

Cells were grown in a minimal seawater medium. The base medium used was from Amarnath et al. (2021)^43^. To reproduce the osmolarity of sea water, it contained 340mM NaCl, 15mM MgCl2, 6.75mM KCl and1mM CaCl2. To reproduce the pH in the ocean, it was buffered using 40mM HEPES salt at pH 8.2. The nitrogen source was 10mM ammonium chloride, the phosphorous source 100mM sodium phosphate and the sulfur source 1mM sodium sulfate. In addition, the medium was supplemented with a mix of trace metals (see Amarnath et al. for full list). Most importantly it contained 1mg/L of iron which was chelated with 4mM tricine. This medium was supplemented with 0.2% w/v chitin flakes (Sigma C9213) or the corresponding carbon source as indicated. Depending on the culture volume, 50mL conical tubes from VWR (89038-658) (for 10mL cultures) or 500mL glass flasks (for 100mL cultures) were used to minimize chitin particles sticking to the walls. Cultures were incubated in water bath shakers at 27C and shaken at 250 rpm.

### Fractionation of the culture

Many of our experiments relied on our ability to separate the various components of the culture as shown in Fig. 1. These components are planktonic cells, particles which contain both cells and excreted proteins and the supernatant which contains excreted proteins.

#### Particle-plankton separation

To separate the particles from the planktonic phase of the culture, samples were taken from the culture with large bore tips (Thermo Scientific 21-236-1A), and spun at a low speed for a short amount of time to sediment the particles.

For OD, RNA, protein as well as microscopy measurements, which required small volumes, samples were transferred to 2mL Eppendorf tubes, spun at 500rpm for 30 seconds on a tabletop centrifuge (Spectrafuge 24D).

For detachment rate, and dialysis experiments, which required larger volumes, 10mL sample were transferred to 15mL conical tubes and spun on a large tabletop centrifuge at 500rpm for 1 minute (Thermo Sorvall Legend T).

In both cases, the planktonic phase was then carefully pipetted out and separated from the remaining particles for further processing.

#### Supernatant extraction

To isolate the supernatant, samples of our culture were passed through a 0.22 um pore vacuum filter (SteriCup Quick Release 150mL S2GPU02RE). To concentrate the excreted proteins, aliquoted samples of the supernatant were spun in 3kDa concentrators (Amicon-Ultra C7715) in a refrigerated tabletop centrifuge at 4k rpm for an hour. They were then consolidated and spun again with the same settings until ∼100x concentration in volume was achieved. The extracted and concentrated samples were kept at 4C until use, but no longer than a day.

### Planktonic OD

500uL samples were taken from the cultures using P1000 pipette tips to make sure chitin particles didn’t clog the tips and transferred to 2mL Eppendorf tubes. Tubes were centrifuged at 500rpm for 30 seconds on a tabletop centrifuge to sediment the chitin particles. 200ul of the planktonic phase were used to determine the planktonic OD of the culture at 600nm using a Genesys 30 spectrophotometer.

### Microscopy

200μl of samples were taken from exponentially growing chitin cultures and fractionated into particles or planktonic cells if needed. Chitin flakes were stained with 1 *μg*/*mL* FITC-WGA and cells were stained with a membrane dye FM4-64 with a concentration of 5*μg*/*mL* . The culture was then fixed with phosphate buffered glutaraldehyde and transferred to a chitin chamber for imaging. The chitin chamber was assembled by appending a precut double-sided 3M tape to a cover glass and then sealed with a cover slip. Chambers were imaged using a Leica TCS SP8 inverted confocal microscope. The WGA signal was read in the GFP channel which was excited with a 488nm diode laser and the FM 4-64 was read in the mCherry channel and excited with a 580 nm diode laser. Fluorescence for both channels was detected through a 40x/1.3 objective and a highly sensitive HyD SP GaAsP detector.

### Dialysis bag experiment

The planktonic and particle components of an exponentially growing chitin culture were separated as described above. In brief 10mL of culture were transferred to a 15mL conical tube (VWR) and subjected to a low speed spin (500rpm) for one minute. The planktonic component was separated from the particles, which were then resuspended in 10mL of chitin-free media. The OD of the resuspended particles was measured and the particles were transferred to a 14kDa MWCO dialysis tube (Sigma D0405) secured with clips. The tube was then inserted in a 500mL glass flask which was filled with the planktonic component. The flask was incubated in a water bath at 27C shaking at 250rpm. The OD outside of the dialysis bag was determined periodically. However, the planktonic OD inside of the bag could only be determined at the end of the experiment upon the reopening of the clips securing the tubing.

### Detachment rate

From exponentially growing chitin cultures from which we note the planktonic OD, we separated the planktonic component from the particles using the method described above. The collected particles were then resuspended in the same volume of fresh, chitin-free media. We then monitored the planktonic OD of this resulting culture at regular time intervals. The detachment rate was obtained from the slope of the linear fit of the resulting data.

### ^32^P incorporation^44^

^32^P-orthophosphate (Perkin Elmer) was added to exponentially growing chitin cultures at 3uCi/ml, and labelling was carried for 2-3 generations before collecting samples. 200 μL samples of the total culture or of only the planktonic fraction were pelleted, washed once with the media buffer and frozen. They were subsequently thawed and resuspended in 2 mL of scintillation cocktail (Liquiscint, National Diagnostics). Radiolabel incorporation was estimated as counts per minute using a Beckmann LS6500 Scintillation counter.

### Protein measurements

Method for the biuret assay adapted from Herbert et al. (1971)^45^. 1.5 mL of an exponentially growing culture of either the full samples, planktonic samples only or particles only was pelleted, washed, resuspended in 200ul of carbonless media buffer, fast-frozen on dry ice and stored. For measurements of extra-cellular protein amounts, 200ul of the concentrated supernatant (see method above) was used. Samples were thawed and the protein concentration measured using the biuret assay. In brief, 0.1mL of 3M NaOH was added to the cell pellet and samples were incubated at a 100C on a heat block for 5 minutes to lyse the cells and hydrolyze the proteins. Samples were then cooled at room temperature for 5 minutes. The biuret reactions are carried out by adding 0.1 ml 1.6% CuSO4 to the samples with thorough mixing at RT for 5 min. Samples were then centrifuged and the absorbance at 555 nm was measured using a spectrophotometer. The same reaction was applied to a series of BSA standards of concentrations ranging from 0-5mg/mL and a standard curve established. Protein amounts in the experimental samples were determined using this standard curve.

### RNA measurement

This method was adapted from You et al. (2013)^46^ and Benthin et al. (1991)^47^ with modifications. For the chitin condition, these experiments were performed in 100mL culture volume shaking in 500mL glass flasks. 1.5 mL of an exponentially growing culture of either the full samples, planktonic samples only or particles only was pelleted, fast-frozen on dry ice and stored. Pellets were thawed and washed twice with 0.7M cold HClO4 then digested for 60 minutes at 37 degrees using 300ul of 0.3M KOH. Samples were periodically stirred. The cell extracts were then neutralized with 100 ul of 3M HClO4 and centrifuged at 13k rpm for 3 minutes. The soluble fraction was collected and the remaining pellets washed twice with 550 ul of 0.5M HClO4. The resulting final volume of 1.5mL was centrifuged once more to eliminate remaining debris and its absorbance at 260 nm was measured using a Bio-Rad spectrophotometer. The RNA concentration was determined as OD260*31/OD600 where the conversion factor is based on RNA’s extinction coefficient.

### RNA synthesis rate measurement

RNA synthesis rate measurement was performed as described in Balakrishnan et al.^48^ (2021) with the following modifications.

To measure RNA synthesis rate of the full culture, 100μl cultures were dispensed into six microfuge tubes from a well-suspended chitin culture. To each tube, 5uCi of 3H-uridine (Perkin Elmer) was added (t=0) and labelling was allowed to continue in the different tubes for different durations (30s, 60s, 90s, 120s, 150s and 180s). Labelling was stopped by adding 100ul boiling lysis buffer (0.1M NaCl, 0.01M Tris, 0.02M EDTA, 0.5% SDS). Each sample was boiled for 2 minutes, chilled on ice for 15 minutes and then RNA was precipitated using an equal volume of ice cold 10% Trichloroacetic acid (TCA).

To measure RNA synthesis rate from planktonic cultures, 1ml of a well-suspended chitin culture was spun down at low speed (500 rpm, 30s) and 975ul of the planktonic culture was moved to a fresh microfuge tube containing 5uCi of ^3^H-uridine (the moment of addition denoted as t=0). At various time intervals (1min, 2min, 3min, 4min, 5min and 6min) 100ul samples were moved to boiling lysis buffer and the samples were boiled, chilled and precipitated as described above for the particles. All precipitated samples were added to 2 mL of scintillation cocktail (Liquiscint, National Diagnostics) and label incorporation was measured using a Beckmann LS6500 Scintillation counter.

### Enzymatic activity

To assay the *in situ* enzymatic activity of the chitinases, 10mL samples were taken at various points along the steady-state growth curve on chitin. 500uL of chloroform were added to permeabilize the cells and prevent their GlcNAc uptake. The tubes were shaken at 250rpm in a 27C water bath shaker.

To assay the enzymatic activity of samples extracted from the supernatant. Equal volumes of the extract were incubated with chitin particles suspended in 10mL of the culture buffer shaking at 250rpm in a 27C water bath.

To monitor the enzymatic activity in both cases, 200ul were sampled from the tubes at regular time intervals and filtered through a 0.22um nylon filter centrifuge tube (Corning Costar Spin-X Centrifuge Tubes). The GlcNAc concentration was then measured from the supernatant using HPLC in the method outlined below.

### HPLC

The HPLC method was adapted form Cremer 2017. In brief, 80µl of filtered sample was then transferred to HPLC sample tubes and analyzed using a Shimadzu Prominence HPLC using RID detection. The HPLC setup was as follows: isocratic HPLC was used with 10mM H_2_ SO_4_ as mobile phase at 0.4ml/min pump speed; samples were kept at room temperature in the autosampler (Shimadzu SIL-10AF); 20µl of sample was injected; samples were separated using ion exchange chromatography; the column (Phenomenex, Rezex ROAOrganic Acid H+ (8%), LC column 300 x 7.8mm) that was kept in a column oven (Shimadzu STO-20A) at 40°C; data from the RID detector (Shimadzu RID-20A) was recorded for 40min. With these settings, the elution time of GlcNAc was between 19.8 and 20.5 minutes. Data was subsequently analyzed with a custom script in MATLAB. In short, peaks of interest were isolated and their baseline adjusted by linearly interpolating the initial and final intensities of the peak. The area under the corrected peaks was then computed. The concentrations corresponding to given areas under peaks were determined by running standards with known concentrations of the compounds of interest.

### SDS-PAGE

Samples (planktonic cells, particles and extra-cellular proteins) were collected from a steady-state chitin culture using the fractionation method outlined above. Cells were lysed and protein reduced by mixing 25μl of sample to 4.75 μl Laemmelli buffer (Bio-Rad #161-0737) and 0.25ul of beta-mercaptoethanol (Bio-Rad #161-0710) and boiling the reaction at 100C for 5 minutes. 30ul of samples were loaded in each well of the pre-cast 10% TGx Mini Protean gels (Bio-Rad #4568033). The ladder used was 1ul of Biorad’s Precision Plus Dual Color #1610374 with bands ranging from 250-10kDa and two bands which fluoresce in UV light at 75 and 25kDa. 1L of running buffer (Bio-Rad #161-0732) was added to the cell and the gels were ran for 40 minutes at 200V. The stain-free fluorescent gels were activated with UV light for 5 minutes and then imaged under a UV gel box. Individual bands of interest were excised using a scalpel that was cleaned upon each incision and stored at 4C for further processing with mass spectrometry.

### Proteomic mass spectrometry

This method was adapted from Hui et. al (2015)^38^. Each sample contained an ^15^N labelled-reference, which allowed to compare unlabeled experimental proteins across growth conditions of interest. Each experimental sample in a series is mixed in an equal amount with a known labelled standard sample as reference, and the relative change of protein expression in the experimental sample is obtained for each protein.

#### Sample collection

For each planktonic culture, 1.5 mL of cell culture at 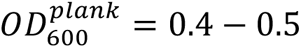 was collected by centrifugation. For chitin cultures, the particles were separated from the planktonic fraction before pelleting. The cell pellet was resuspended in 0.2 mL of water and fast frozen on dry ice. For the extra-cellular sample, 200ul of the 100x concentrate of the supernatant (see extraction method above) was used.

#### Sample preparation

^15^N labelling was achieved by using ^15^N labelled ammonia as the sole nitrogen source. A balanced mixture of the two ^15^N labelled cell samples (from a glycolytic fast condition on glucose and a gluconeogenic slow growth condition on acetate) was prepared as a universal reference. These conditions were chosen such as to obtain a wide coverage of the proteome without biasing a particular growth condition. 50 μg of the reference was mixed with 50 μg of each experimental sample. This balanced preparation (equal amounts of total protein) enabled the measurement of proteome mass fraction for each protein.

Proteins were precipitated by adding 100% (w/v) trichloroacetic acid (TCA) to a final concentration of 25%. Samples were left to stand on ice for a minimum of 1 h. The protein precipitates were spun down by centrifugation at 13,200g for 15 min at 4 °C. The supernatant was removed and the pellets were washed with cold acetone and dried in a Speed-Vac concentrator.

The pellets were dissolved in 80 μl 100 mM NH_4_ HCO_3_ with 5% acetonitrile (ACN). 8 μl of 50 mM dithiothreitol (DTT) was added to reduce the disulfide bonds before the samples were incubated at 65 °C for 10 min. Cysteine residues were modified by adding 8 μl of 100 mM iodoacetamide (IAA) followed by incubation at 30 °C for 30 min in the dark. Proteolytic digestion was carried out by adding 8 μl of 0.1 μg μl−1 trypsin (Sigma-Aldrich) with incubation overnight at 37 °C. The peptide solutions were cleaned by using PepClean C-18 spin columns (Pierce, Rockford, IL). After drying in a Speed-Vac concentrator, the peptides were dissolved into 10 μl sample buffer (5% ACN and 0.1% formic acid). For samples containing chitin particles, only the soluble portion of the pellet was used.

#### Data acquisition

Data acquisition was adapted from Hui et al. (2015)^38^ and Dai et al. (2016)^49^. MS data was acquired using an AB Sciex 5600 TripleTOF spectrometer with the injection of 2 µg tryptic peptides, with MS1 accumulation time of 250 ms and MS2 accumulations of 150 ms.

#### Protein identification

Protein identification was adapted from from Hui et al. (2015)^38^ and Dai et al. (2016)^49^. In brief, raw data files (.wiff and .wiff.scan formats) were converted to profile and centroided mzML formats. Centroided mzML files were converted to mzXML using tools included in the Trans-Protemic Pipeline (TPP) and searched using X!Tandem (https://thegpm.org) against the UniProt *Vibrio sp.* 1A01 database (organism ID 314742) supplemented with common protein contaminants, enzymes and reversed peptide decoy sequences. The peptide-spectrum match tolerances were set at 50 ppm and 100 ppm for the precursor and product ions, respectively. The TPP tools PeptideProphet and iProphet were used to score the peptide–spectrum matches and the search results were combined into a consensus library using SpectraST.

#### Relative protein quantification

Relative protein quantitation was adapted from Hui et al. (2015)^38^ and Dai et al. (2016)^49^. Using an in-house quantification software, Massacre, on the consensus library, we quantified the relative intensities of the ^14^N (light) peptides to ^15^N (heavy) peptides. In brief, the intensity for each peptide is integrated over a patch in RT, m/z space that encloses the envelope for the light and heavy peaks. After collapsing data in the RT dimension, the light and heavy peaks are fit to a multinomial distribution (a function of the chemical formula of each peptide) using a least squares Fourier transform convolution routine, which yields the relative intensity of the light and heavy species. The ratio of the unlabeled to labelled peak intensity is obtained for each peptide in each sample. A confidence measure for each fit is calculated from a support vector machine (SVM) trained on a large set of user scoring events.

#### Absolute protein quantification

The absolute protein level for each sample was obtained by dividing the ^14^N spectral counts for each protein by the total spectral counts detected in each sample in the ^14^N channel.

## DATA AVAILABILITY

The mass spectrometry proteomics data have been deposited to the ProteomeXchange Consortium (Vizcaíno et al, 2014) via the UCSD MassIVE partner repository with the dataset identifier PXD034003 (Username: MSV000089499_reviewer, Password: vibriosp1a01). Summary tables including both absolute and relative quantitation as well as functional grouping can be found in Tables S2 and S3. All other data that support the findings of this study are available from the corresponding author upon request

All correspondences and requests for materials should be directed to Terry Hwa

## ACKNOWLEDGEMENTS

We thank Julia Schwartzman and Otto Cordero for providing strains and helpful discussions. We are very grateful to Farooq Azam for general discussions which were extremely informative. We are also grateful to members of the Hwa research group for their helpful input and feedback. Illustrations were created using biorender.com. This work is supported by the Simons Foundation through the Principles of Microbial Ecosystems (PriME) collaboration (Grant no. 542387 to TH, and 542395 to JRW.). Some figures were created using BioRender.com.

## AUTHOR CONTRIBUTIONS

T.H and G.G conceived the study, designed experiments and wrote the manuscript. G.G conducted all experiments with assistance from R.B with the radioactivity. V.P processed proteomics samples and data and T.C contributed to modeling work.

## EXTENDED DATA FIGURES AND TABLES

**Extended Data Figure 1:**
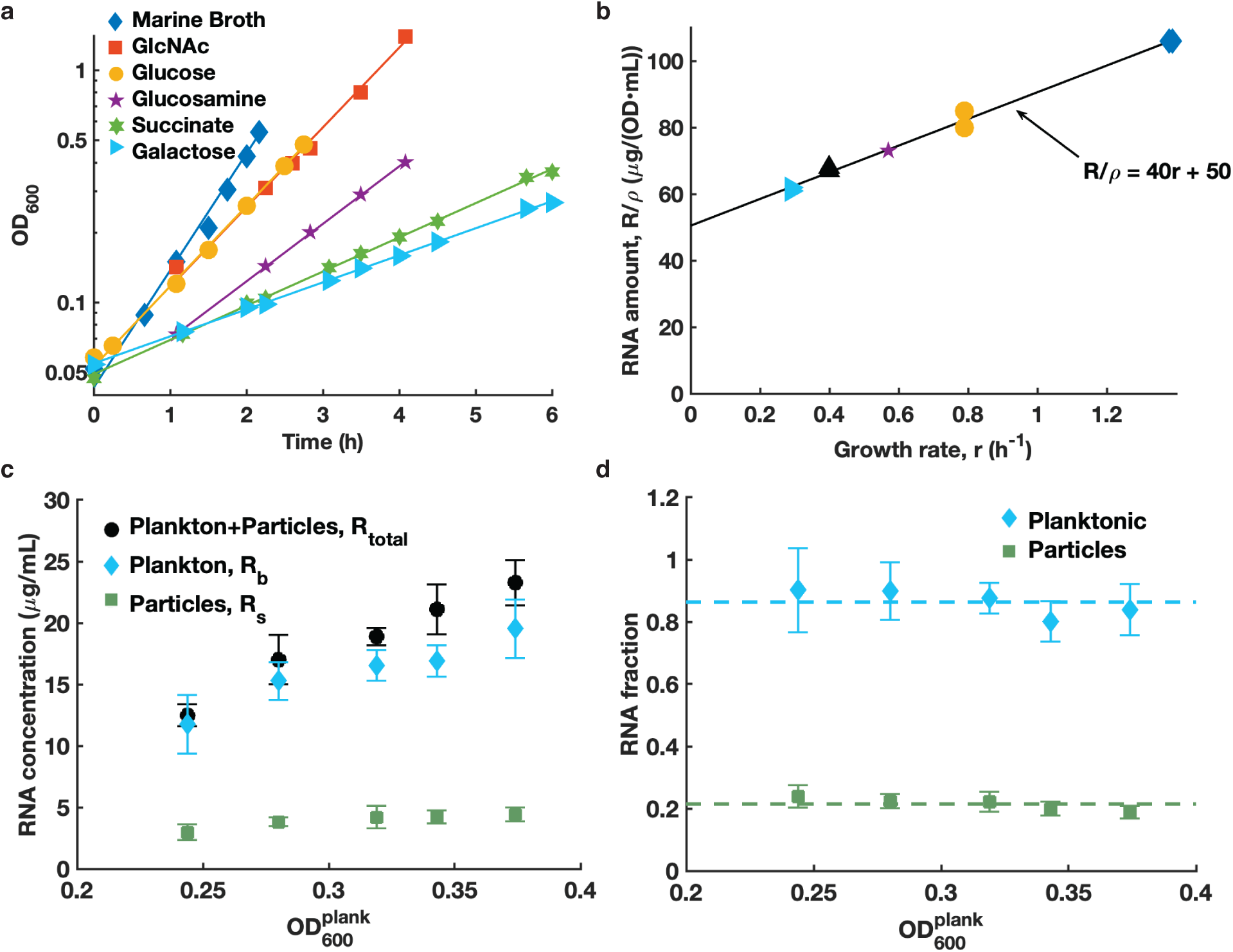
Quantifying the RNA abundance of cells grown in different conditions. **a)** Growth curves of 1A01 grown in minimal media with different carbon sources. Cultures were inoculated from exponentially growing pre-cultures with the same carbon source. The replication rates (*r*) were obtained from the exponential fits and ranged from 1.15 ℎ^-1^ to 0.26 ℎ^-1^; see Table S2 for values. **b)** RNA amounts per OD•mL of culture, *R*/*ρ*, was obtained for exponentially growing cultures in different carbon sources and plotted against the respective replication rates, *r* (same symbols as Panel a). The RNA content exhibits a linear dependence on the replication rate *r* as indicated by the line of best-fit given in the plot. In the main text, we determined that the replication rate of particle-associated cells and planktonic cells are respectively *r*_*s*_ ≈ 0.26ℎ^-1^ and *r*_*b*_ ≈ 0. The best-fit line then allows us to deduce *R*_*s*_: *ρ*_*s*_ ≈ 60 *μg*/(*OD* • *mL*) and *R*_*b*_: *ρ*_*b*_ ≈ 50*μg*/(*OD* • *mL*) for these two subpopulations of cells. Together with the result *R*_*b*_: *R*_*total*_ ≈ 0.72 from Fig. 1d, we obtain *ρ*_*b*_: *ρ*_*total*_ ≈ 0.75 ± 0.06. **c)** Direct measurement of RNA content of the different subpopulations of the chitin culture. RNA concentration was measured for samples of the full culture, planktonic cells and on particles only using perchloric acid precipitation. We observe that the planktonic samples (blue diamonds) were consistently below the full samples (black circles) and similarly to Fig. 1c, we interpret this difference as the biomass accumulating on the surface of the particles. Moreover, the linear increase of RNA concentration with OD in the three samples indicates again that the culture has reached a steady-state. The error bars correspond to the standard deviation across three biological replicates. **d)** To determine the planktonic fraction of the culture (blue diamonds), we took the ratio of the blue diamonds to the black circles in Panel c. This fraction remained constant throughout growth, further establishing that the chitin culture was in a steady-state. The data provides another estimate of the fraction of RNA in the planktonic subpopulation, with *R*_*b*_: *R*_*total*_ ≈ 0.82 ± 0.1, which is comparable to the estimate obtained in Fig. 1d using radio-labeling. Using the RNA content described in Panel b, the fraction of the planktonic subpopulation is estimated to be *ρ*_*b*_: *ρ*_*total*_ ≈ 0.8 ± 0.1, consistent with the estimate of *ρ*_*b*_: *ρ*_*total*_ ≈ 0.75 ± 0.06, given above.

**Extended Data Figure 2:**
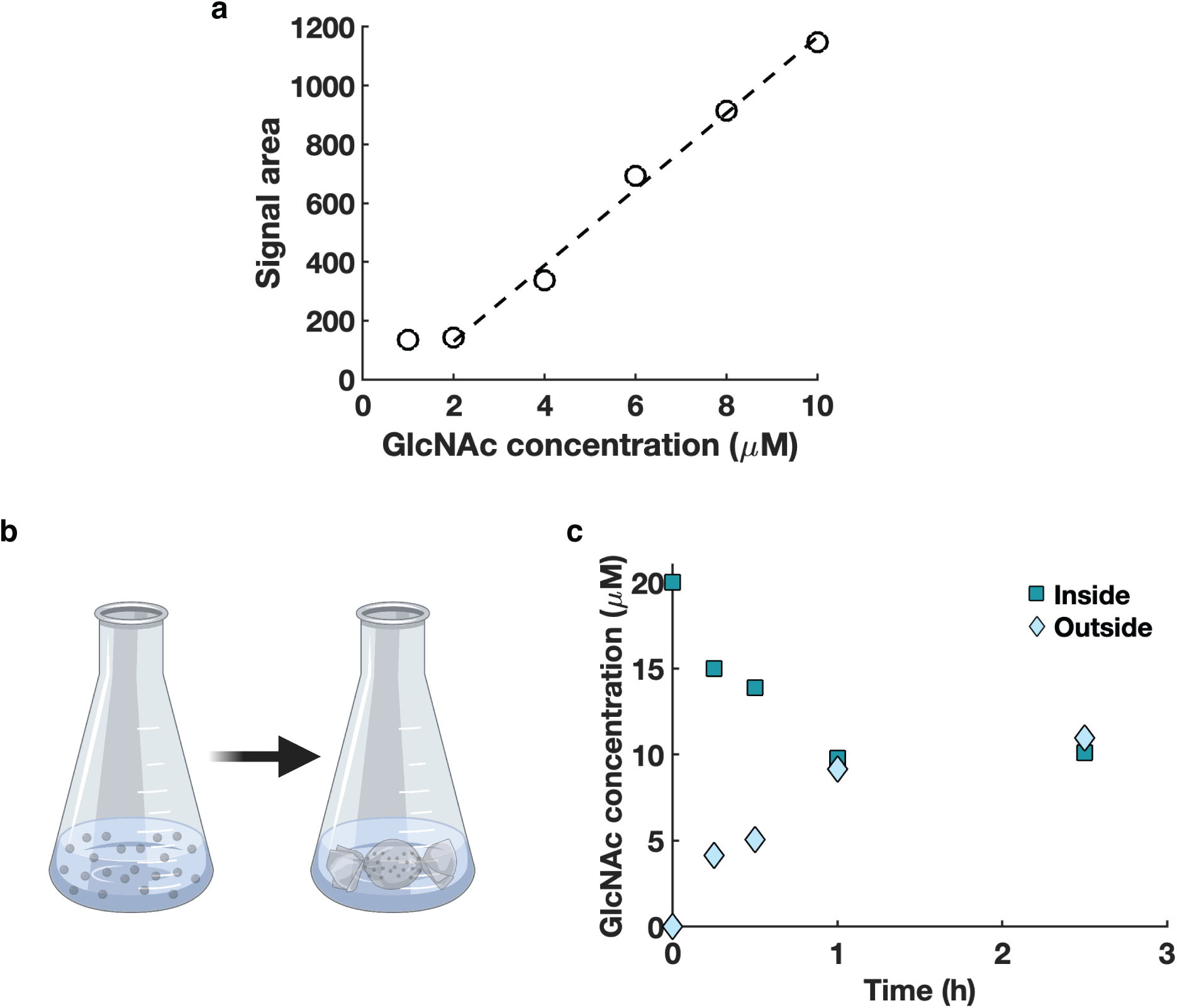
Detecting the dynamics of GlcNAc in the media. **a)** Standard curve showing the detection limit of GlcNAc using HPLC. Several high dilutions of GlcNAc, ranging from 10 − 2 *μM* were prepared in the minimal media. The samples were passed through a hydrophobic column equipped with a refractive index detector (see HPLC in Methods). Peaks corresponding to GlcNAc eluted between 19.8 and 20.5 minutes, and the total signal area under the curve was determined. The signal area scaled linearly with GlcNAc concentration at the input all the way down to 2 *μM*, below which peaks were not detected. Analyzing the supernatant of a growing chitin culture with the same method, yielded an absence of measurable peak, indicated the GlcNAc concentration in the supernatant is below 2 *μ*, the detection limit of the HPLC. **b)** Illustration of the dialysis setup in Figs. 2c-d. Chitin particles were separated from planktonic cells in the middle of exponential growth. Chitin particles were resuspended in the same volume of fresh media without particles and placed inside a dialysis bag with a 14kDa molecular weight cutoff. This pore size means that small molecules such as GlcNAc can exchange but not enzymes, cells nor chitin particles. The dialysis bag was inserted into a flask and the planktonic cells were transferred to the flask, to the exterior of the bag, with equal volumes inside and outside of the dialysis bag. Outside of the dialysis bag, the OD was tracked regularly over a time period of 30 hours, while inside of the bag only the initial and final ODs were recorded due to the difficulty of handling the dialysis tubing. **c)** 10mL of cell-less media with 20 *μM* of GlcNAc was placed in a dialysis bag and incubated with the same volume of media without GlcNAc outside of the bag. The concentrations start equilibrating immediately after the start of the experiment, and reach their equilibrium value by 1h. This indicates that with the full chitin culture in Figs. 2c-d, a low concentration of GlcNAc generated inside the bag can freely exchange between the interior and exterior of the bag if not taken up by cells inside the bag.

**Extended Data Figure 3:**
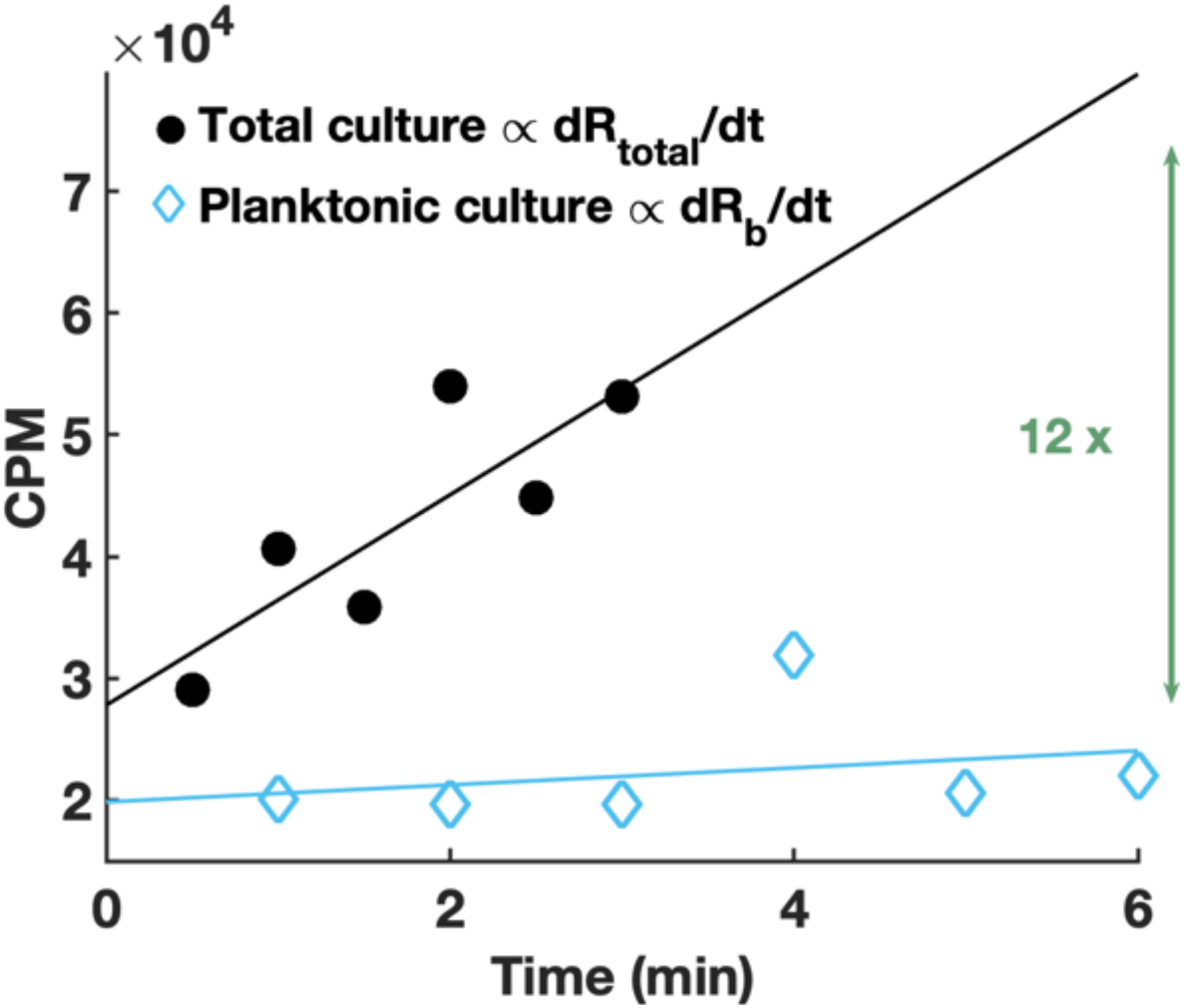
Instantaneous RNA synthesis rate measurements in the chitin culture. From an exponentially increasing chitin culture at 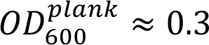, samples of the full culture (including chitin particles and planktonic cells) were compared to samples containing planktonic cells only. The instantaneous RNA synthesis rate, a proxy for the cells’ metabolic rate, was measured in both samples by pulsing ^3^H-Uridine and immediately tracking its incorporation into the total culture and planktonic culture (see see RNA synthesis rate in Methods). The plot shows the radioactivity reading as a function of time. The solid lines are linear best fits to the data and their slope represents the metabolic rate in each sample. Specifically let *A*_*total*_ = *r*_*s*_*ρ*_*s*_ + *r*_*b*_*ρ*_*b*_ be the metabolic rate of the full culture and *A*_*b*_ = *r*_*b*_*ρ*_*b*_ be the metabolic rate of the planktonic fraction. The result *A*_*total*_/*A*_*b*_ ≈ 12 indicates that the particle-associated cells were the main contributor to RNA synthesis (and hence cell replication) even though they comprised a minority of the biomass as was determined in Fig. 1 and Extended Data Fig.1. The replication rates on and off the particles can be compared by taking the ratio *A*_*total*_/*A*_*b*_ and expressing *r*_*b*_ given its contribution to the total biomass: *r*_*b*_ = (*r*_*s*_ (*ρ*_*total*_/*ρ*_*b*_ − 1))/(*A*_*total*_/*A*_*b*_ − 1) = *r*_*s*_/33. Given the estimate of *r*_*s*_ = 0.26ℎ^−1^ (main text), this indicates that *r*_*b*_ ≈ 0.008ℎ^-1^ ≪ *λ*, making *r*_*b*_ practically negligible.

**Extended Data Figure 4:**
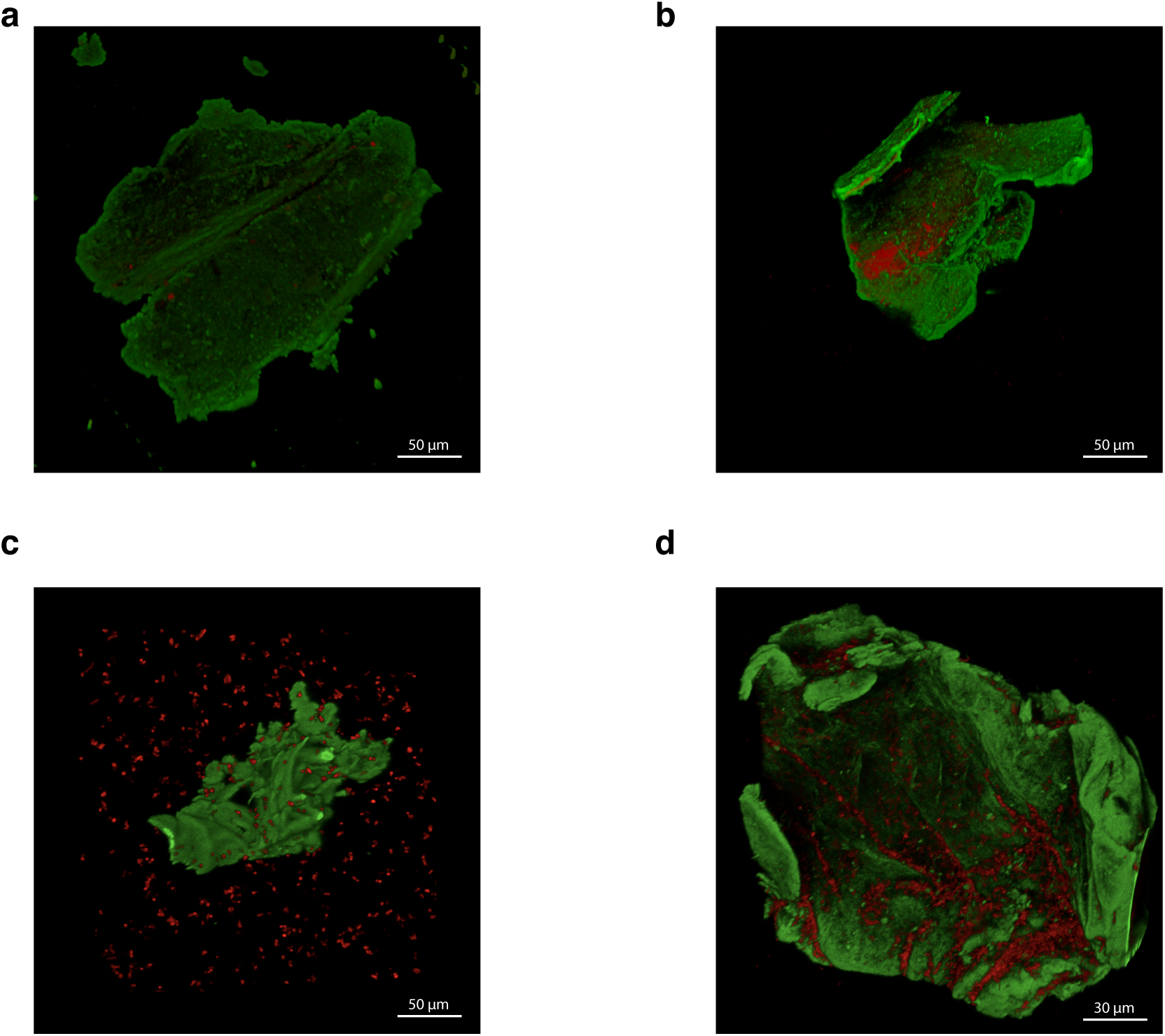
Microscopy images of chitin particles from growing cultures. Samples were incubated with two fluorescent dyes: FITC-WGA, here shown in the green channel and FM 4-64, a cell membrane dye, shown in the red channel, and fixed with glutaraldehyde. Imaging of the samples was done in custom-built chambers (see microscopy in Methods) under a confocal microscope with 40x magnification. **a)** Sterile chitin flake incubated with both dyes. FITC-WGA specifically binds to chitin particles, though some FM 4-64 can also be seen in the background. **b)** Chitin particles were isolated from an exponentially increasing chitin culture at 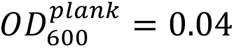 and resuspended in fresh media. Red (membrane dyed) cells can be seen to bind to chitin and form microcolonies. **c)** A sample including both planktonic cells and chitin particles from an exponentially increasing chitin culture at 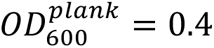. Consistent with bulk measurements, more planktonic cells are observed than surface-associated cells. **d)** From the same growing culture as in Panel c, chitin particles were isolated and resuspended in fresh media to remove planktonic cells. In both c) and d), the surface of particles is not saturated with cells even at this moderately high OD. This suggests that cell detachment from particles is not a result of a “space limitation” on the particle surface.

**Extended Data Figure 5:**
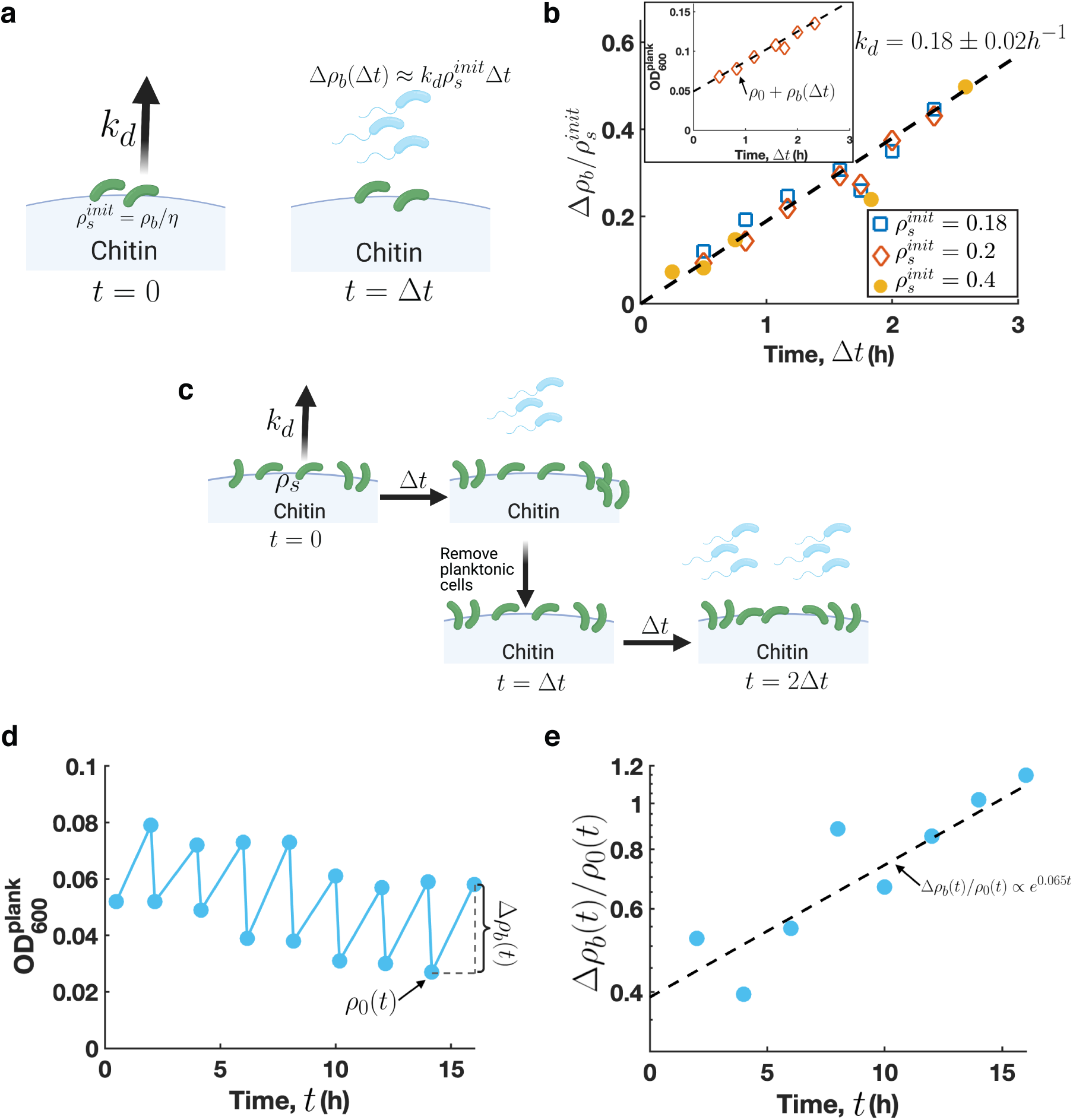
1A01 has a dispersal lifestyle. **a)** Chitin particles pre-colonized with cells (green cells) were isolated at various time (*τ*) from an exponentially increasing chitin culture, and resuspended in the same volume of fresh media without chitin particles. The amount of particle-associated cells was estimated using the planktonic OD at the time of particle isolation *ρ*(*r*) and the constant planktonic fraction *ρ*_*b*_: *ρ*_*total*_ obtained in Fig. 1, i.e., 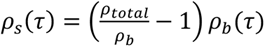. This is taken as the initial cell density 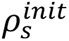 of the resuspended culture. For each resuspended culture, the planktonic OD (*ρ*_*b*_, giving the density of blue cells) was measured at regular time intervals *Δt*. The increase in planktonic OD is expected to be linear in time with a rate *k*_*d*_ for some time after resuspension. **b)** Inset: the planktonic OD for one such resuspensions increases linearly starting from a background reading *ρ*_5_, due to the turbidity caused by small chitin particles. The dashed line is the line of best fit to the data with a slope 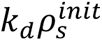. Main: Traces of the planktonic OD (*Δρ*_*b*_) were normalized to the initial amount of cells on the particles 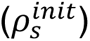 and plotted as a function of time. The normalized traces collapse on top of one another. The slope of the line of best fit (dashed line) allows an estimate of the detachment rate, with *k*_*d*_ ≈ 0.18 ± 0.02 ℎ^-1^, with the error given by the 95% confidence interval of the fit. **c)** Similarly to Panel a, pre-colonized chitin particles from a 10mL exponentially increasing chitin culture were separated and re-suspended into the same volume of fresh media without chitin particles. After a time interval of Δ*t* = 2ℎ, the planktonic OD increased by an amount Δ*ρ*_*b*_ as a result of cell detachment and with the rate observed Panel b. At this point, the particles were sedimented, and resuspended in the same volume of fresh media without chitin particles. This process, which effectively removed all the planktonic cells and thereby prevented reattachment (effectively setting the attachment rate *k*_*a*_ ≈ 0), was repeated 8 times, for a total of 16 hours. **d)** Traces of the planktonic OD as a function of time for successive resuspension cycles. From an initial OD measurement *ρ*_5_, which is the background OD due to the residual turbidity of small chitin particles, we observed an increase in the planktonic OD, Δ*ρ*_*b*_, after the 2h time cycle. The background OD exhibits a time dependence *ρ*_5_(*t*) due to the successive removal of small particles in the fractionation process. The data shows that in the absence of planktonic cells, the particle-associated cells alone were able to sustain replication and shedding of new planktonic cells in the duration of our experiments, demonstrating that the planktonic culture is not necessary for the replication of cells on the particles. **e)** From the traces in Panel b, we plot the planktonic increase relative to the background OD as a function of time. The normalization by the dynamic background OD allows to adjust for the loss of small particles and hence of surface-associated cells from the culture during the sedimentation process. The dashed line is obtained by fitting an exponential model to the data, with the growth exponent being 0.065 ℎ^-1^. Since the cell detachment rate is proportional to the number of cells on particles, the relative increase in planktonic OD, Δ*ρ*_*b*_(*t*)/*ρ*_5_(*t*), is thus a proxy for the increase in cell density on the particles. We see that this relative increase in planktonic OD is exponential, and that its rate matches the exponential rate measured in an undisturbed chitin culture, *λ* ≈ 0.06 ℎ^-1^ (Fig. 1). Since the rate of exponential increase in the planktonic population is hardly affected by the removal of planktonic cells from the chitin culture, we conclude from the data that the attachment rate *k*_*a*_ ≪ *λρ*_*s*_/*ρ*_*b*_ = 0.02ℎ^-1^ is thus negligible compared to other rates in our culture.

**Extended Data Figure 6:**
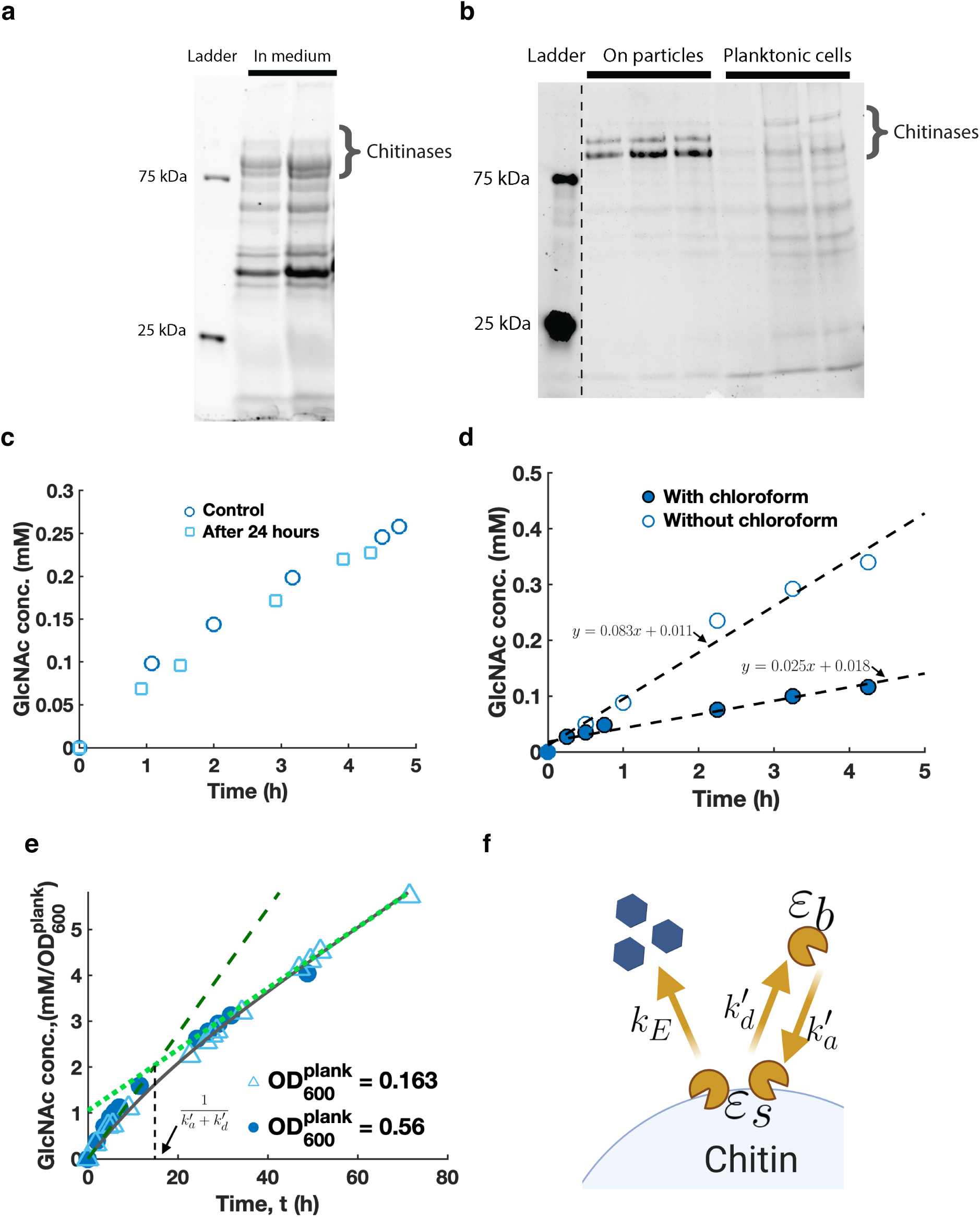
Chitinase activity and properties. **a-b)** SDS-PAGE gels were run to separate and identify differences in the proteomes of the three fractions of the chitin culture (planktonic cells, particles, and medium after filtering out planktonic cells). To identify the proteins in each band, the bands were excised, solubilized and analyzed using mass spectrometry for identification; see SDS-PAGE in Methods. Bands corresponding to chitinases (with molecular weight ∼100kDa) are indicated on the images. Gel image: light intensity and contrast were adjusted. Original gel images are available upon request. **a)** Samples were collected from a steady-state chitin culture at 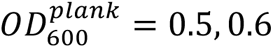 (from left to right). 100 mL of culture was passed through a 0.22 *μm* filter, and the filtrate was concentrated 100-fold using 3kDa concentrators (see fractionation in Methods). The resulting protein mixture was loaded onto the gel. Chitinases are found in the supernatant. See ED Table 1 for the amount quantified. Other unrelated lanes on the same gel were cropped out of the image shown. **b)** Throughout a steady-state chitin culture at multiple planktonic ODs, (at 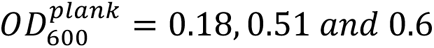 respectively from the left to the right lane), samples were collected and particles separated from planktonic cells. Particles were concentrated 4-fold compared to the volume of planktonic cells to make sure loading amounts were in the same range. After boiling and reducing the proteins (see SDS-PAGE in Methods), they were loaded onto the gel. From the band intensities, it is clear visually that the chitinases are enriched on the particles. This observation is quantitatively confirmed by mass spectrometry, which yields a 5-fold enrichment of chitinases per cell on the particles compared to planktonic cells (see ED Table 1). Gel image: three unrelated lanes between the ladder and the particle samples shown were cropped out from the original image. **c-d)** Proteins from the supernatant of a steady-state chitin culture were collected at 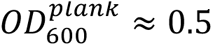 and concentrated using the method described above. They were then incubated with fresh chitin flakes and the activity of the enzymes was determined by measuring the GlcNAc concentration in the supernatant at different times using HPLC. **c)** To test the stability of the enzymes, they were resuspended in the same buffer used in our chitin culture. The activity of the enzymes was assayed for similar amounts immediately after their collection (circles) and after being incubated in the same shaking conditions as our culture at 27C for 24 hours (squares). This procedure resulted in the same rate of GlcNAc accumulation indicating that the enzymes were stable for the duration assayed. **d)** Because we used chloroform as a way of disabling nutrient uptake in our *in-situ* measurements of enzymatic activity (Figure 4 and Panel e), we independently tested the effect of chloroform on the absolute activity of the chitinases by incubating similar amounts of supernatant enzymes with chitin particles in our culture buffer with and without the addition of chloroform. For each case, we measured the concentration of GlcNAc in the media at regular time intervals (filled and open circles, respectively). The dashed lines represent the linear curves of best fit for the data and the best-fit parameters are indicated on the plots. The ratio of the slopes of the two lines was about 3.3, indicating that the activity of the chitinases enzymes was reduced ∼3.3 fold due to chloroform treatment. **e)** Similarly to Fig. 3c, we tracked the accumulation of GlcNAc after treating exponentially growing chitin cultures with chloroform to inhibit carbon uptake by cells in the culture. In this case, GlcNAc accumulation was followed over a longer timescale (3 days) to assess possible changes in chitinase activity, through observing changes in the rate of GlcNAc accumulation. Samples were collected at planktonic ODs of 0.16 (open triangles) and 0.63 (filled circles). The resulting GlcNAc accumulation traces were normalized to the planktonic OD at sampling and we found that the traces collapsed onto each other after normalizing by the initial OD. The reaction was kept at 27C and shaking throughout the course of our measurement. The longer timescale of this experiment showed that after 15 hours, there was a change of rate in the accumulation of the GlcNAc concentration (compare dashed to dotted green lines). The second rate was thereafter maintained for ∼3 days. This is incompatible with a gradual degradation of chitinases but rather suggests that the system finds a new equilibrium due to the exchange dynamics (attachment and detachment) of the enzymes. **d)** To interpret the data in Panel e) and extract the value of the relevant parameters, we formulate the following model governing the GlcNAc accumulation rate. surface-attached enzymes (*ε*_*s*_) which produce GlcNAc with a catalytic rate *k*_*E*_ as in Fig. 3 are exchanged with the bulk enzymes (*ε*_*b*_) with attachment and detachment rates respectively 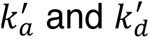. We plot the best-fit solution for the traces as the solid black line. The full solution of this model can be found in Supp. Note II. Here we briefly summarize the results: Initially, as explained in Figures 4b-d, the rate of GlcNAc accumulation is proportional to the enzyme’s catalytic rate *s*_1_ = *κ*_*E*_*m*_*E*_*ε* . At longer times the slope decreases and corresponds to the equilibration of attached and detached enzymes, with 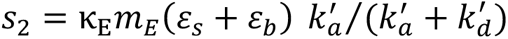. The ratio of the two slopes is therefore: 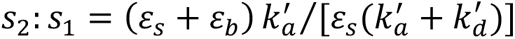. The change from one regime to the other occurs at a point 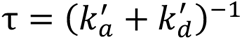. Thus, with the empirical values of the quantities *s*_2_: *s*_1_ and τ obtained from the data in panel a), as well as the ratio *ε*_*b*_: *ε*_*s*_ obtained from ED Table 1 (ratio of the third to the first entry in the last row), we can determine the attachment and detachment rates 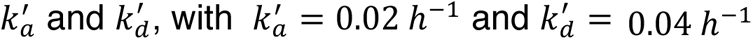

**Extended Data Figure 7:**
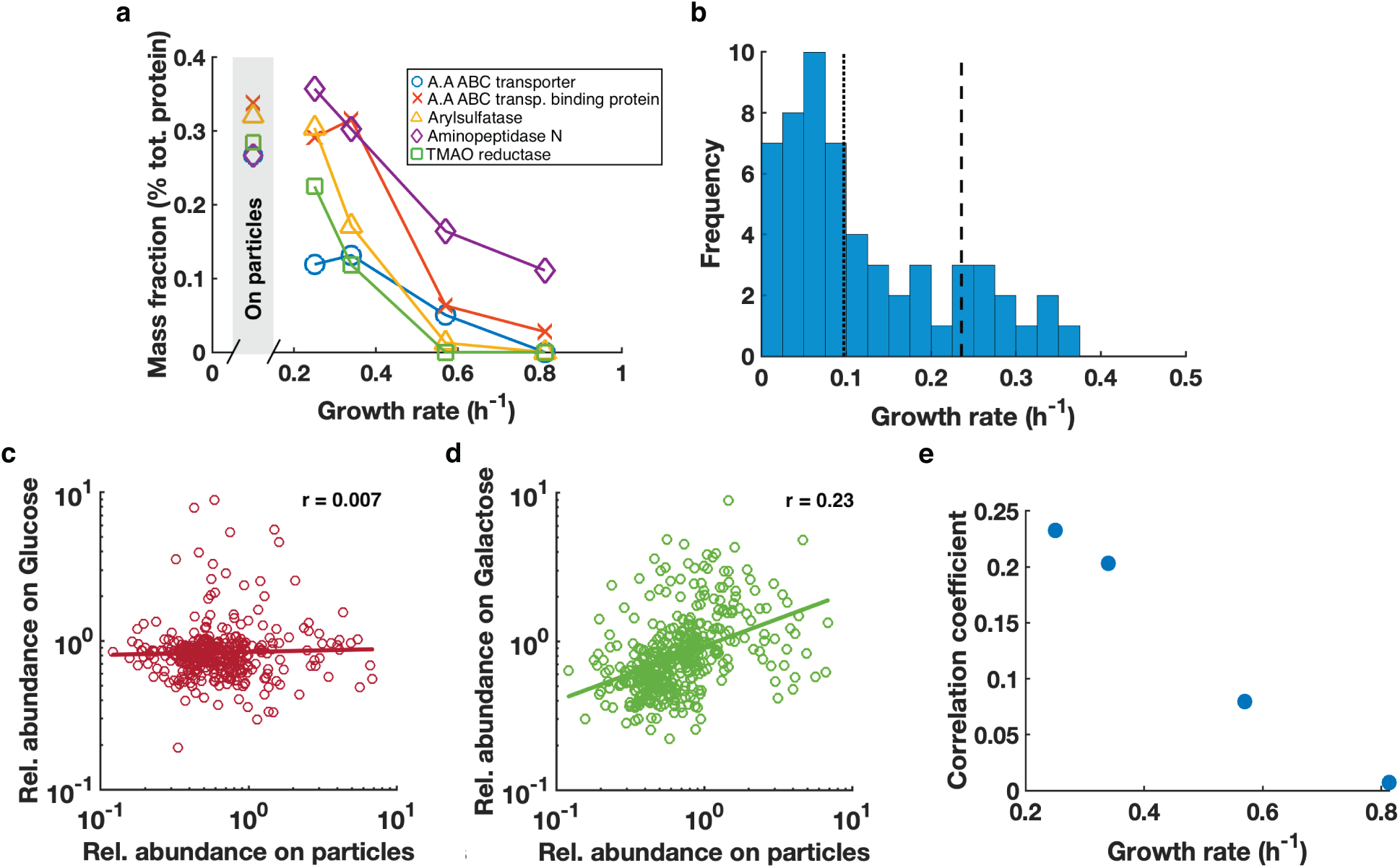
Analysis of proteomic data. **a)** Specific examples of proteins whose abundances increased strongly on carbon sources resulting in poor growth. The mass fraction is obtained as the fractional spectral count of these individual proteins in each sample. The x-axis represents the growth rate of the culture under carbon sources of varying quality: Glucose (0.81/h), GlcNAc (0.8/h), Glucosamine (0.57/h), Succinate (0.34/h) and Galactose (0.25/h). The data in the shaded bar show the mass fraction of these proteins on chitin particles (excluding the chitinases). The abundance of these proteins is more comparable to that on slow growth in poor carbon sources than on fast growth. **b)** Analysis of individual ribosomal proteins. For each ribosomal protein, a linear curve was fitted for its abundance as a function of growth rate using the series described in panel A with varying carbon quality. This linear fit was used as a predictor of the growth rate of surface-associated cells depending on the abundance of individual ribosomal proteins on the particles subtracted of chitinases. The histogram (blue bars) shows the result of the predicted growth rates of surface-associated cells based on each of the 53 ribosomal proteins detected. They yield an average predicted growth rate of 0.23 ℎ^-1^ (dashed vertical line) with the median being even lower, 0.1 /h, (dotted vertical line). **c-d)** Scatter plots of the relative abundances of proteins from surface-associated cells and cells grown on **c)** glucose and **d)** galactose. Each data point represents the abundance of a protein relative to our ^15^N labelled standard in both conditions examined. Our standard was composed of a mixture of cells extracted from a fast (glucose) and a medium (succinate) growing condition and injected in equal amount into all of our samples to allow for comparisons between them (see media recipe in Methods). Stronger correlation (0.23) is seen between the proteome of surface-associated cells and cells grown on galactose (poor carbon source giving growth rate of 0.25/h), while weaker correlation (0.007) is seen between surface-associated cells and cells grown on glucose (good carbon source giving growth rate of 0.81/h). **e)** The correlation coefficient is calculated between the proteome of cells grown in each of the carbon sources studied and that of particle-associated cells as done in panels c and d except for GlcNAc to avoid biases due to the substrate’s nature. Plotting the correlation coefficient with the growth rate of the corresponding carbon source, we see that higher correlation is progressively obtained for slower carbon sources, suggesting that particle-associated cells are carbon-limited.

**Extended Data Figure 8:**
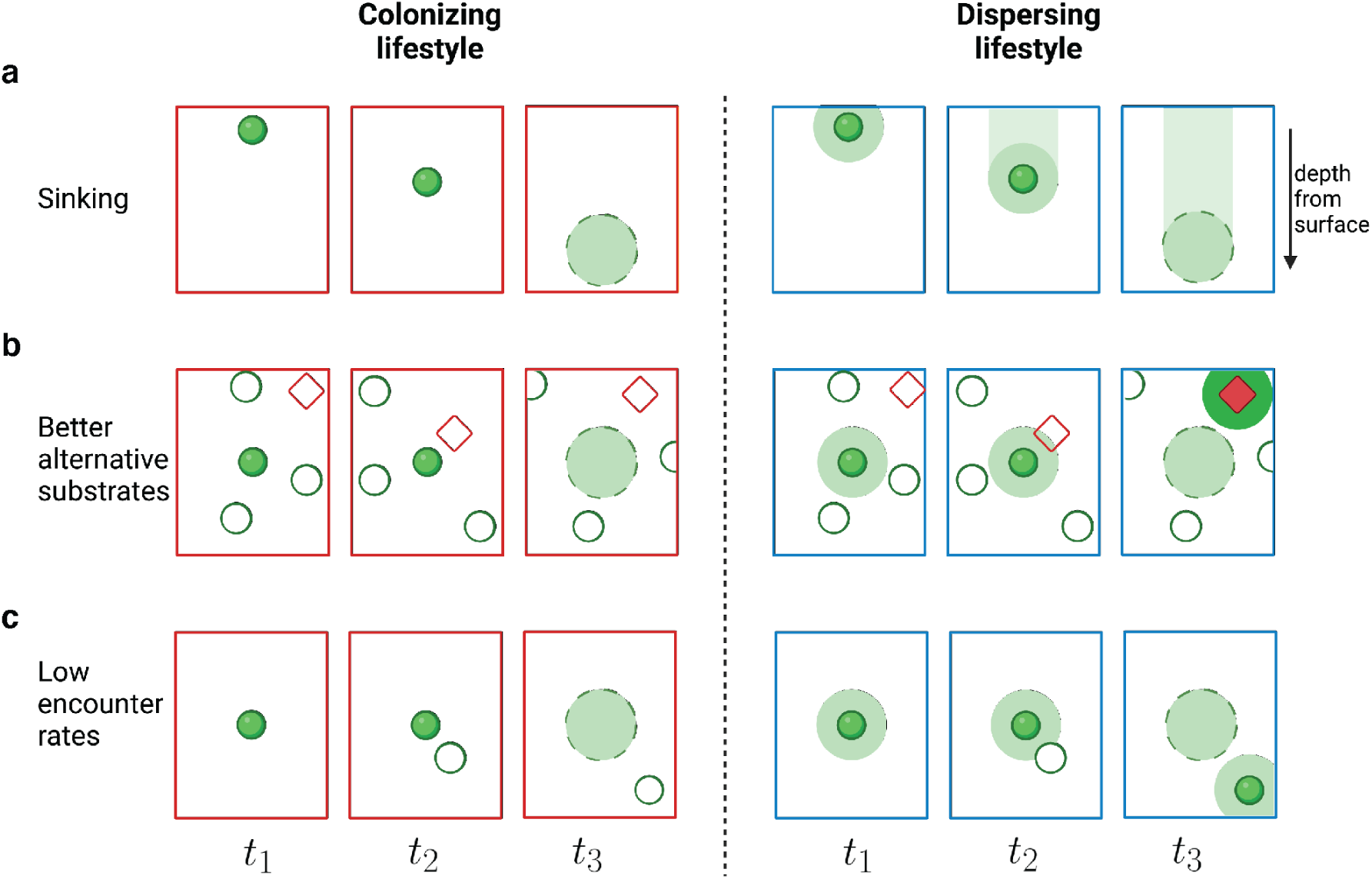
Possible environmental factors favoring or disfavoring dispersal. Diverse scenarios are presented favoring either the colonizing or the dispersing lifestyle. Briefly, in the colonizing lifestyle, cells do not detach from particles until they are fully consumed, after which all cells are released in a “burst”. In this case, the “bursted” population would all die unless one of the cells colonized a fresh particle during the cell lifetime following the burst. In the dispersing lifestyle, a fraction of the population is continuously shed as the particle is being degraded. This allows for cells to seed fresh particles at any time before the particle is fully consumed. For each scenario, the frames from left to right illustrate possible dynamics of the population on and surrounding the particle being degraded. Filled symbols represent colonized particles and the green shade represents planktonic cells. Open symbols represent fresh particles. Diamonds represent a different solid substrate supporting faster growth. **a)** The vertical direction represents depth from the water surface. High detachment rates allow the population to maintain itself along the water column and avoid inhospitable depths where oxygen and other factors may become limiting for growth. **b)** A high detachment rate allows the seeding of a particle made of a different substrate (red diamond) providing faster growth. This can be beneficial for certain *Vibrio sp*. species that favor, e.g., alginate^50, 51^. However, we remark that if detachment reduces the fitness of the population on the existing niche as commonly believed, then exploring alternative resource patches would be more costly. **c)** In situations where the particle density in the environment is low and thus the encounter rate is also low, a high dispersal rate increases the probability of encountering a fresh particle during the “lifetime” of a colonized particle. If seeding itself is not difficult, this increases the probability of encounter for dispersing cells. On the other hand, if successful seeding is a low probability event while encountering fresh particles is not, then “bursting” would be more advantageous given the larger number of cells released at burst. This may arise from the requirement on the absolute numbers of cells for the successful colonization of new particles^2,52^, or due to Allee effects arising from other mechanisms.

**Extended Data Figure 9:**
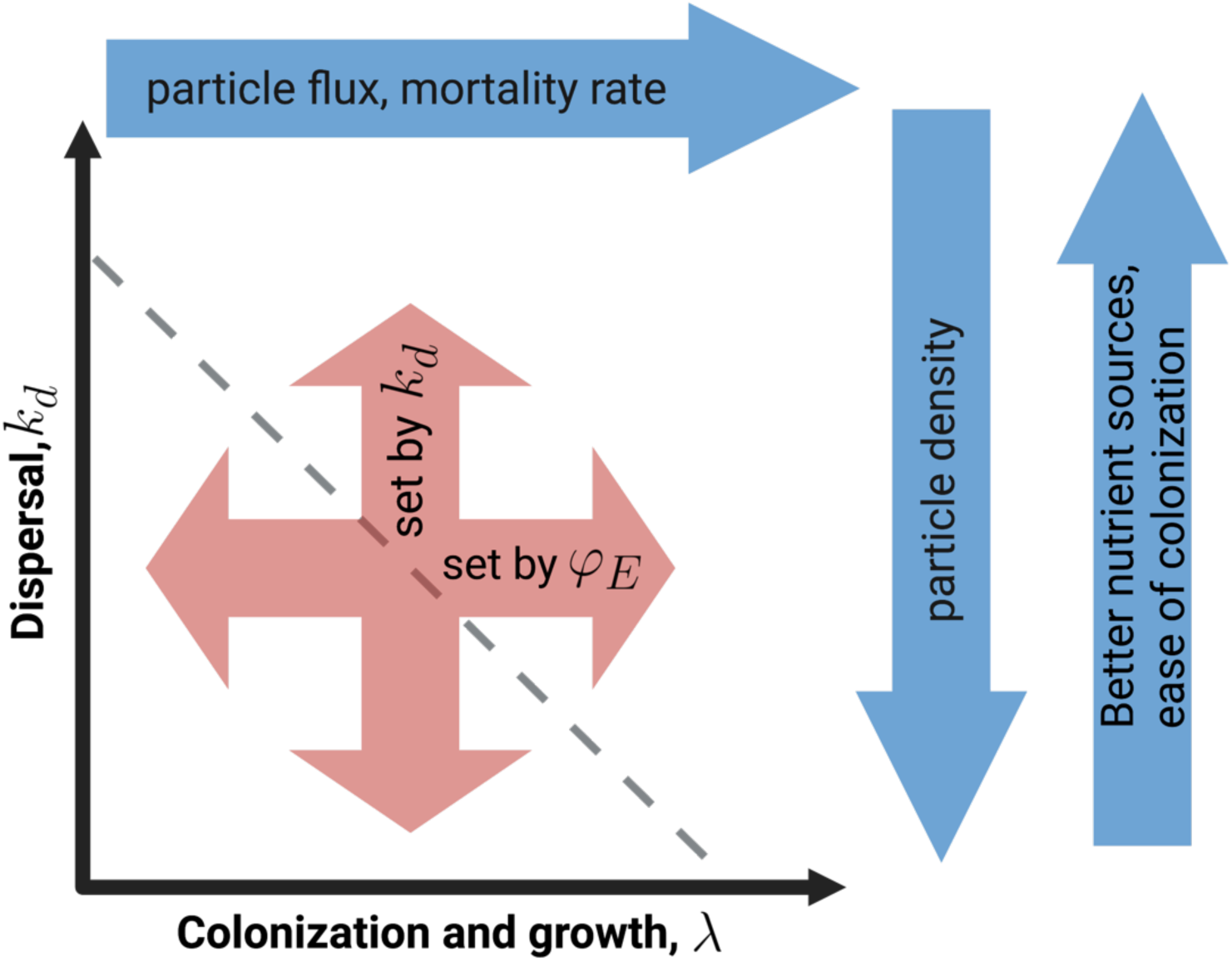
Cellular and environmental factors determining growth and dispersal. For bacteria like *Vibrio sp. 1A01* which release their secreted chitinases, the rate of increase of the population colonized on chitin particles, *λ*, is decoupled from the rate of detachment from the particles, *k*_*d*_, circumventing the commonly held trade-off between growth and dispersal, illustrated by the dashed grey line. Instead, growth and dispersal can be separately set molecularly (red arrows), with the population increase rate *λ* set by *φ*_*E*_, the allocation towards chitinase synthesis (Eq. (6)). The two molecular parameters *k*_*d*_ and *φ*_*E*_ are in turn dependent on environmental factors such that the population-level fitness, which involves both growth and successful colonization of many particles, increases over long timescales. We speculate on a number of such environmental factors, represented by the blue arrows. Examples of factors favoring detachment are illustrated in Extended Data Fig. 8. Factors setting the rate of increase of the colonized population may include the rate of particle influx and the rate of mortality on particles (due to grazing, phage killing…) since the survival of the population requires it to grow above the rate of mortality, and as much as allowed by the overall nutrient influx, but not above it to avoid long periods of starvation. We remark that even with growth and dispersal decoupled, an anti-correlation between these two traits may arise if environmental factors favoring one factor also disfavors another. e.g., a low particle density resulting from low particle influx would favor slow growth and high detachment.

**Extended Data Table 1:**
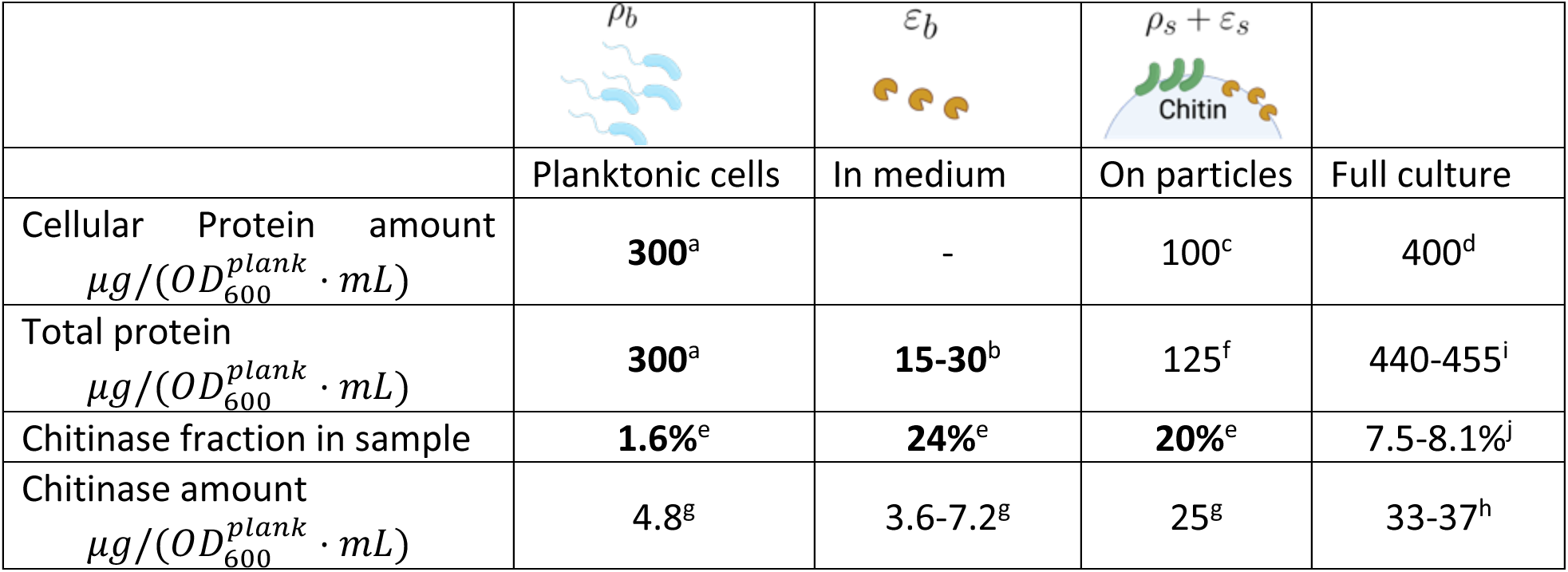
Chitinase abundance in different components of the chitin culture. Planktonic cells and particles were separated using the sedimentation method described in Figure 1a and the supernatant was obtained after passage of the entire culture through a 0.2 *μm* pore size filter and subsequent concentration through a 3kDa concentrator. See details in Methods. Numbers in bold were directly measured while plain numbers were computed. a- The protein amount in planktonic cells *M*_*ρb*_ was measured using the biuret assay. b- The protein amount in the medium was determined by the biuret assay in the concentrated supernatant (see protein measurement in Methods). c- The cellular protein amount on particles *M*_*ρs*_ was computed using the ratio *η_b_* = 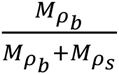 found in Fig. 1, with 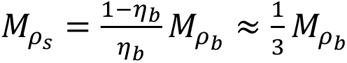 d- The total cellular protein amount is *M*_*ρb*_ + *M*_*ρs*_ e- The chitinase fraction was determined as the sum of chitinase spectral counts (called A0A2G1AS61, A0A2G1AX49 and A0A2G1AVJ7 in the UniProt database, see Table S3) divided by the total spectral counts in each sample. We refer to this fraction on the particles as *f*_*s*_, in planktonic cells as *f*_*b*_ and in the medium as *f*_*m*_. In particular, *f*_*s*_ = *M*_*ε*_*s*__/(*M*_*ρs*_ + *M*_*ε*_*s*__). f- The total amount of proteins (including cellular proteins and chitinases) on the particles 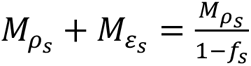 g- The chitinase amount in each column is computed as the product of the entries of the second and third row in each column. h- The total chitinase amount is the sum of left 3 entries of the last row. i- The total protein amount in the entire chitin culture *M*_*total*_ is the sum of the left 3 entries of 2^nd^ row). a) The chitinase fraction in the total culture is the ratio of the entries of the 4^th^ and 2^nd^ row of the last column.

| | $\rho_b$ | $\varepsilon_b$ | $\rho_s + \varepsilon_s$ | |
| --- | --- | --- | --- | --- |
|  | Planktonic cells | In medium | On particles | Full culture |
| Cellular Protein amount<br>$\mu g / (OD_{600}^{plank} \cdot mL)$ | **300<sup>a</sup>** | - | 100<sup>c</sup> | 400<sup>d</sup> |
| Total protein<br>$\mu g / (OD_{600}^{plank} \cdot mL)$ | **300<sup>a</sup>** | **15-30<sup>b</sup>** | 125<sup>f</sup> | 440-455<sup>i</sup> |
| Chitinase fraction in sample | **1.6%<sup>e</sup>** | **24%<sup>e</sup>** | **20%<sup>e</sup>** | 7.5-8.1%<sup>j</sup> |
| Chitinase amount<br>$\mu g / (OD_{600}^{plank} \cdot mL)$ | 4.8<sup>g</sup> | 3.6-7.2<sup>g</sup> | 25<sup>g</sup> | 33-37<sup>h</sup> |

