## Supplementary Information for "Sustained growth and rapid dispersal of chitin-associated marine bacteria"

Guessous, G. et al.


#### SUPPLEMENTARY TABLES 2

**Table S1: Summary of parameters and their values.** 3
**Table S2: Proteomics data** Excel table attached
**Table S3: Summary of proteomics data by functional group** Excel table attached

#### SUPPLEMENTARY NOTES 4

**I- A population-level model for chitin degradation** 4
1- General population-level model 4
2- Case of secreted chitinases 6
3- Case of cell-bound enzymes 11
4- A conflict between privatization and high detachment 13
5- On the attachment rate 14

**II- On chitinase dynamics** 18

**III- Spatial model for growth on a chitin particle** 20
1- Case with a constant planktonic cell density 20
2- Case with a spatially-dependent planktonic cell density 22

#### SUPPLEMENTARY REFERENCES 25

### SUPPLEMENTARY TABLE

| Parameter | Unit | Notation | Value | Source |
|---|---|---|---|---|
| Max. replication rate on GlcNAc | $h^{-1}$ | $r_{max}$ | $0.8 \pm 0.05$ | Fig. 1b |
| Population growth rate on chitin | $h^{-1}$ | $\lambda$ | $0.06 \pm 0.01$ | Fig. 1b |
| Density of planktonic cells | $OD_{600}^{plank} \approx 10^9\ cells/mL$ | $\rho_b$ | various | Supp. Ref. [1] |
| Fraction of planktonic cells | -- | $\eta_b$ | $0.75 \pm 0.06$ | Fig. 1c |
| Number of particle-associated cells per culture volume | $OD_{600}^{plank}$ equivalent | $\rho_s$ | various | $\rho_b \cdot (1-\eta_b)/\eta_b$ |
| Rate of cell detach. from chitin | $h^{-1}$ | $k_d$ | $0.18 \pm 0.02$ | Ext. Data Fig. 5b |
| Cell Yield of GlcNAc | $OD_{600}^{plank}/mM$ | $Y$ | 0.16 | Supp. Ref. [1] |
| Cell dry mass | $\mu g/ml/OD_{600}^{plank}$ | $m_{cell}$ | $575 \pm 25$ | Supp. Ref. [2] |
| Cell protein mass | $\mu g/ml/OD_{600}^{plank}$ | $m_P$ | $300 \pm 30$ | Ext. Data Table 1 |
| Molecular weight of chitinase | $kDa$ | $m_E$ | 100 | Ext. Data Fig. 6 |
| Protein yield of GlcNAc | μg/mL/mM | $Y_P$ | $48 \pm 5$ | $Y \cdot m_P$ |
| Biomass yield of GlcNAc | μg/mL/mM | $Y_b$ | $92 \pm 4$ | $Y \cdot m_{cell}$ |
| Total cellular protein on particles | $\mu g/ml/OD_{600}^{plank}$ | $\rho_s m_P/\rho_b$ | 100 | $m_P(1-\eta_b)/\eta_b$ |
| Mass fraction of chitinase amongst all proteins on particle | % | $f_s$ | 20 | Ext. Data Table 1 |
| Mass fraction of chitinase on particles among all chitinases | % | $f_E$ | $71 \pm 5$ | footnote a |
| Chitinase amount on particle | $\mu g/ml/OD_{600}^{plank}$ | $m_E \varepsilon_s/\rho_b$ | 25 | $\frac{\rho_s m_P}{\rho_b} \cdot \frac{f_s}{1-f_s}$ |
| GlcNAc production flux | $mM/h/OD_{600}^{plank}$ | $J_n/\rho_b$ | 0.6 | footnote b |
| Specific chitinase activity | nmol/μg/h | $\kappa_E$ | $24 \pm 5$ | Fig. 3 |
| Fraction of biomass production towards chitinases on particles | % | $\varphi_E$ | $3 \pm 0.2$ | footnote c |
| Fraction of total proteome that are chitinases | % | $\chi_E$ | $4.9 \pm 0.5$ | footnote d |
| Fraction of biomass production towards all chitinases | % | $\varphi_{E,tot}$ | $4.2 \pm 0.3$ | $\varphi_E / f_E$ |
| Max. allocation for cell growth | % | $\varphi_{max}$ | $34 \pm 4$ | footnote e |

**Table S1: Summary of parameters and their values.** We summarize the key cellular and molecular quantities obtained in this work, by either direct measurement (indicated by the Figure or Table where the result is derived), or indirectly from measured quantities (indicated by the relation used to compute them).

a) $f_E$ is the ratio of the amount of chitinase found on particle to the total amount of chitinase in the culture; values of the latter are given as the 3rd and 4th entries of the last row of Extended Data Table 1.

b) We can check that the GlcNAc production flux indeed supports the growth of the entire culture such that $J_n Y_b = \lambda\big(m_{cell}(\rho_b + \rho_s) + m_E(\varepsilon_b + \varepsilon_s)\big) \approx \lambda(m_{cell}(\rho_b + \rho_s))$. Solving for the growth rate, we find $\lambda = \frac{J_n}{\rho_b}\eta_b Y_b = 0.07h^{-1}$, quantitatively corresponding to the observed growth rate.

c) $\varphi_E$ is defined as the fraction of the total nutrient flux that goes towards chitinase synthesis, i.e., $\varphi_E \equiv \frac{J_E}{J_{rep}+J_E} = \frac{m_E\varepsilon_s}{(\rho_b+\rho_s)m_{cell}+m_E\varepsilon_s}$. It can be rewritten as $\varphi_E = \frac{m_E\varepsilon_s/\rho_b}{m_{cell}\cdot(\rho_b+\rho_s)/\rho_b+m_E\varepsilon_s/\rho_b}$ and determined using the parameter values listed above.

d) $\chi_E$ is defined as the fraction of proteins in the culture that are chitinases such that $\chi_E \equiv m_E\varepsilon_s/\big((\rho_s + \rho_b)\cdot m_P + m_E\varepsilon_s\big)$. It can be related to the nutrient flux fraction $\varphi_E$ from $\chi_E = \varphi_E/[b + (1 - b)\varphi_E]$ where $b = m_P/m_{cell}$.

e) $\varphi_{max}$ is the maximum fraction of the nutrient flux dedicated to the synthesis of proteins necessary for growth in the fastest growth condition. We first estimated $\chi_{max}$ by observing that at the fastest growth rate $\chi_{max} \approx 50\%$ (since the translational apparatus is 18%, enzyme biosynthesis 17%, transporters 5%, glycolytic enzymes 7% and TCA enzymes 4% (see Summary sheet in Table S3)). We next converted this quantity from a protein fraction to a nutrient flux fraction $\varphi_{max}$ since $\varphi_{max} = b\chi_{max}/[1 - (1 - b)\chi_{max}]$ where $b = m_P/m_{cell}$.

### SUPPLEMENTARY NOTES

#### I-A population-level model for chitin degradation

In this section we present the most general framework for describing the processes that lead to chitin degradation at the population level, including both cell and enzyme dynamics. After describing this framework, we will examine the properties of two specific bacterial strategies: one in which enzymes are secreted and released extra-cellularly and another in which enzymes remain attached to the cell surface.

##### 1- <u>General population-level model</u>

*Vibrio sp.* 1A01 cells are grown in a minimal medium with chitin flakes as their sole source of Carbon and Nitrogen. We refer to planktonic or **b**ulk components with the subscript $b$ and to **s**urface associated components with the subscript $s$.

The total volume of the culture is $V$ and the amount of chitin in the culture is described by the weight density $\phi_w$. Given chitin's density $d_c = 1.4\ g/mL$, the number density of chitin particles is $\phi_v = \phi_w/d_c$. For a chitin particle of volume $v_p$, the number of chitin particles in the culture is: $N_c = \phi_v\ V/v_p$. Hence the number density of chitin particles is $\rho_c = N_c/V = \phi_v/v_p$. The particle volume $v_p$ depends on the shape of the particles, which we approximate as spheres of radius $R_0$ such that $v_p = 4/3\pi R_0^3$.

We define cellular and enzymatic variables in the chitin culture:

- Let $N_b$ be the total number of cells in the bulk. $\rho_b = N_b/V$ is the cell density in the bulk.
- Let $N_s$ be the total number of cells associated with the surface of the chitin particles. $\rho_s = N_s/V$ is the surface-associated cell density. We note that these also include cells that are in the immediate vicinity of particles, in the "chitosphere", as we explain more clearly in Supp. Note III. The number of cells per particle is $\rho_s/\rho_c$.
- Let $N_{\varepsilon_b}$ be the number of chitinases in the bulk. $\varepsilon_b = N_{\varepsilon_b}/V$ is the chitinase density in the bulk.
- Let $N_{\varepsilon_s}$ be the number of chitinases associated with the surface of the chitin particles. $\varepsilon_s = N_{\varepsilon_s}/V$ is the chitinase density in the bulk. The number of enzymes per particle is $\varepsilon_s/\rho_c$.
- $n$ is the GlcNAc concentration on the surface of particles.

With these components, the most general model at the population level is:

$$\frac{d\rho_b}{dt} = k_d\rho_s + (r_b - k_a)\rho_b \qquad \text{(S1)}$$

$$\frac{d\rho_s}{dt} = (r_s - k_d)\rho_s + k_a\,\rho_b \qquad \text{(S2)}$$

$$\frac{d\varepsilon_b}{dt} = \beta_b\rho_b + k_d'\varepsilon_s - k_a'\varepsilon_b \qquad \text{(S3)}$$

$$\frac{d\varepsilon_s}{dt} = \beta_s\rho_s - k_d'\varepsilon_s + k_a'\varepsilon_b \qquad \text{(S4)}$$

$$\frac{dn}{dt} = k_E \varepsilon_s - \mu \rho_s - j_D \qquad \text{(S5)}$$

Where:

- $k_d$ and $k_a$ are the cell detachment and attachment rates respectively,
- $r_b$ and $r_s$ are the cell replication rates in the bulk and on the surface of the particles respectively,
- $k'_d$ and $k'_a$ are the enzymes' detachment and attachment rates respectively,
- $\beta_b$ and $\beta_s$ are the chitinase synthesis rates by the bulk and surface-associated cells respectively,
- $\mu$ is the nutrient uptake rate by surface-associated cells,
- $k_E$ is the catalytic rate of the chitinases referred to in the main text as $k_E = \kappa_E m_E$, where $\kappa_E$ is the catalytic rate per enzyme mass and $m_E$ the enzyme mass,
- $j_D$ is the nutrient loss due to diffusion, referred to in the main text as $j_{loss}$

Given this system, let's follow the nutrient flux and define a few flux quantities. Let $J_n$ be the total nutrient generation flux in units of mM/h. To ensure flux balance in this closed system, this flux supports the total biomass growth such that $J_n = J_{rep} + J_E + j_{loss}$. $J_{rep}$ is the nutrient flux dedicated to cellular replication and biomass while $J_E$ is the nutrient flux dedicated to chitinase synthesis (see Fig. 3 in Main text). In section III, we argue why $j_{loss}$ can be neglected from this calculation. This is also supported by the experimental observation that planktonic cells aren't replicating (see Fig. 2 in main text). To relate the molecular variables defined in the system above (Eqs. S1-S5) to these fluxes, we consider $Y_b$ as the biomass yield of 1A01 growing on GlcNAc, which acts as a conversion factor between biomass and nutrient concentration.

To ensure flux balance, the total nutrient flux is thus:

$$J_n Y_b = \Sigma_i \left(\dot{\rho}_i + \dot{\varepsilon}_i\right) \qquad \text{(S6)}$$

And the flux going towards chitinase synthesis only:

$$J_E Y_b = \Sigma_i \, \dot{\varepsilon}_i \qquad \text{(S7)}$$

Similarly to the main text, it is convenient to introduce the quantity $\varphi_E$, which describes the fraction of the total nutrient flux going towards chitinase synthesis:

$$\varphi_E \equiv J_E / (J_{rep} + J_E) \qquad \text{(S8)}$$

This fraction of the flux directed to chitinase synthesis is a key control parameter since it describes the cost incurred to cells as a result of chitinase production. If there were no cost associated with chitinase production, the replication rates $r_i$, would only depend on the nutrient concentration through the Monod form:

$$r_i(n/K) = r_{max} \frac{n/K}{1+n/K}$$

Where:

- $r_{max}$ is the maximal growth rate of 1A01 in minimal media with GlcNAc as the sole carbon source,
- $n/K$ is the nutrient concentration available in units of the Monod constant $K$.

However, given the necessity of synthesizing chitinases and its potential cost on the proteome of 1A01, we modify this form of the replication rate by referring back to results in Scott et al. (2010)[3]. To complete the description of the system, we introduce the proteomic cost of chitinase synthesis on the replication rate of surface associated cells. For a fixed nutrient level, the replication rate can be expressed as follows:

$$r_i(n/K, \varphi_E) = r_{max} \frac{n/K}{1+n/K} \cdot (1 - \varphi_E/\varphi_{max}) \qquad \text{(S9)}$$

Where:

- $\varphi_{max}$ is defined as the maximum nutrient flux fraction dedicated to growth (refer to Figure 4B in Scott et al. (2010)).

To intuitively understand the form of this function, we consider two limiting cases. On the one hand if $\varphi_E = 0$, growth is nutrient limited as it would be in a chemostat, the replication rate solely depends on nutrient levels: $r_s(n/K, \varphi_E = 0) = r_{max} \frac{n/K}{1+n/K}$. On the other hand, for situations in which $\varphi_E \neq 0$, there would be a cost to chitinase production with a maximal effect when $\varphi_E = \varphi_{max}$, at which point the cells can no longer replicate since all of their available biomass is divested from biosynthetic proteins and ribosomes and goes towards chitinase production. In this case: $r_i(n/K, \varphi_E = \varphi_{max}) = 0$.

We note that even though here the control parameter $\varphi_E$ is expressed as a nutrient flux fraction, it can easily be converted to the variables usually used in models including the cost of protein overexpression[4,5]. These models are generally written in terms of proteomic fractions $\chi_i$, since these are the experimentally accessible quantities. In particular, we can write $\chi_i = b\chi/[1-(1-b)\chi]$ where $b = m_P/m_{cell}$ converts between protein mass and biomass of the cell.

#### 2- Case of secreted chitinases

##### *a- Experimental constraints*

Experimental findings allow to constrain the previous model given the measured parameters for the growth *Vibrio sp.* 1A01 on chitin.

In Fig. 1, we demonstrate using several ways that our system is in steady-state, with both the planktonic and particle-associated fraction increasing exponentially at the same exponential rate, $\lambda$. We refer to this parameter as the population increase rate.

In Fig. 2, we demonstrate that planktonic cells aren't replicating, (i.e: $r_b = 0$). Moreover, in Extended Data Table 1 we show that their chitinase production is negligible (i.e: $\beta_b = 0$).

Since the planktonic cells aren't replicating this is a strong experimental indication that the diffusive loss term $j_{loss}$ is negligible compared to the other fluxes. Another way to convince ourselves that $j_{loss}$ is negligible in this steady-state is to compare it to the other terms in Eq. S5. The loss term can be modelled as: $j_{loss} = 4\pi D R_0 \rho_c Sh$ where $D$ is the diffusion coefficient and $Sh$ the Sherwood number. On the surface of the particles and at steady-state, $\varepsilon_s$ and $\rho_s$ increase exponentially while the diffusion term doesn't. A more complete model which shows that the diffusion loss term decays with a finite screening length is solved in Section III of this note.

In Extended Data Fig. 5 we experimentally determine the value of $k_d$ and establish that $k_a$ is small compared to $\lambda$.

These estimates allow to reduce Eq. S1-S5 to this simplified system of equations:

$$\frac{d\rho_b}{dt} = k_d \rho_s \quad \text{(S10)}$$

$$\frac{d\rho_s}{dt} = (r_s - k_d)\rho_s \quad \text{(S11)}$$

$$\frac{d\varepsilon_b}{dt} = k'_d \varepsilon_s - k'_a \varepsilon_b \quad \text{(S12)}$$

$$\frac{d\varepsilon_s}{dt} = \beta_s \rho_s - k'_d \varepsilon_s + k'_a \varepsilon_b \quad \text{(S13)}$$

$$\frac{dn}{dt} = k_E \varepsilon_s - \mu \rho_s \quad \text{(S14)}$$

The total nutrient flux in units of $[mM/h]$, is $J_n = k_E \varepsilon_s$ . Given the system is at steady-state with an exponential rate of increase $\lambda$, we solve Eqs. S10-S14 to obtain the following solution:

$$\eta \equiv \rho_b/\rho_s = k_d/\lambda \quad \text{(S15)}$$

$$\lambda = r_s(n/K, \varphi_E) - k_d \quad \text{(S16)}$$

$$\eta' \equiv \varepsilon_b/\varepsilon_s = k'_d/(\lambda + k'_a) \quad \text{(S17)}$$

$$\beta_s \rho_s = (\lambda + k'_d)\varepsilon_s - k'_a \varepsilon_b \quad \text{(S18)}$$

$$k_E \varepsilon_s = J_n \quad \text{(S19)}$$

In Figure 1, we measure $\lambda$ and $\eta$. These measurements help estimate $r_s$.

Experiments in Figures 4 and Extended Data Fig. 6, help determine the enzyme properties $k_E$, $k'_a$ and $k'_d$ as well as their steady-state repartition $\eta'$. We note that in the main text we treat the simpler case in which enzymes automatically attach to particles and don't detach from them. This corresponds to the case where $k'_d = 0$ and $\varepsilon_b = 0$.

Using the definitions in Eqs. S6-S7, as well as the steady-state solution, we express the various nutrient fluxes as:

$$J_n Y_b = r_s \rho_s + \beta_s \rho_s \quad \text{(S20)}$$

$$= \lambda(\rho_b + \rho_s + \varepsilon_b + \varepsilon_s) \quad \text{(S21)}$$

And

$$J_E Y_b = \beta_s \rho_s \quad \text{(S22)}$$

$$= \lambda(\varepsilon_b + \varepsilon_s) \quad \text{(S23)}$$

Which leads to the steady-state expression for $\varphi_E$ using its definition in Eq.S8:

$$\varphi_E = \frac{\varepsilon_b+\varepsilon_s}{\rho_b+\rho_s+\varepsilon_b+\varepsilon_s} \qquad \text{(S24)}$$

The two key control parameters that cells can control within the timescale of our experiment (i.e: as a result of gene regulation) are $\varphi_E$ and $k_d$. On the other hand, enzyme parameters such as $k_E$, $k_d'$ and $k_a'$ are dictated by the genetic sequences available to 1A01 and can only change on longer time scales and as a result of sequence evolution. Below, we examine the effect of changing these two key parameters on the properties of the steady-state solution described here.

*b- <u>Examining the effect of $\varphi_E$ and $k_d$ on the system</u>*

Our goal in this section is to examine the dependence of $\lambda$ on the control parameters $\varphi_E$ and $k_d$.

Subtracting Eqs. S20 and S22 and expressing $J_n$ as in Eq. S19 allows to find the replication rate $r_s$ as a function of the control parameters:

$$Y_b(1-\varphi_E)J_n = r_s\rho_s \qquad \text{(S25)}$$
$$\rightarrow r_s = Y_b(1-\varphi_E)k_E\varepsilon_s/\rho_s \qquad \text{(S26)}$$

Thus, given Eq. S16,

$$\lambda = (1-\varphi_E)\kappa_E m_E Y_b \varepsilon_s/\rho_s - k_d \qquad \text{(S27)}$$

Rearranging this expression for $m_E\varepsilon_s/\rho_s$ and plugging the result into the steady-state expression for $\varphi_E$, in Eq. S20 we get:

$$\varphi_E = \left(\frac{1+\eta'}{1+\eta}\varepsilon_s/\rho_s\right)/\left(1+\frac{1+\eta'}{1+\eta}\varepsilon_s/\rho_s\right) \qquad \text{(S28)}$$

Using Eqs. S15 and S17, $\frac{1+\eta'}{(1+\eta)}\varepsilon_s/\rho_s$ can be written as:

$$\frac{1+\eta'}{(1+\eta)}\varepsilon_s/\rho_s = \frac{\lambda+k_a{}'+k_d{}'}{\lambda+k_a{}'}\frac{\lambda}{(1-\varphi_E)\kappa_E Y_b} \qquad \text{(S29)}$$

Thereby cancelling the $k_d$ dependence of $\varphi_E$:

$$\varphi_E = \frac{\frac{\lambda+k_a{}'+k_d{}'}{\lambda+k_a{}'}\frac{\lambda}{(1-\varphi_E)k_E Y_b}}{1+\frac{\lambda+k_a{}'+k_d{}'}{\lambda+k_a{}'}\frac{\lambda}{(1-\varphi_E)k_E Y_b}}$$

Rearranging for $\lambda$ in the previous equation, we have:

$$\lambda^2 + (k_a' + k_d' - \kappa_E Y_b \varphi_E)\lambda - k_a'\kappa_E Y_b \varphi_E = 0$$

Now solving this quadratic form, we find $\lambda(\varphi_E)$:

$$\rightarrow \lambda(\varphi_E) = \frac{1}{2}\left(k_E Y_b \varphi_E - (k'_a + k'_d) + \sqrt{(k'_a + k'_d - k_E Y_b \varphi_E)^2 + 4k'_a k_E Y_b \varphi_E}\right) \quad \text{(S30)}$$

This analysis indicates that surprisingly the exponential population increase rate, $\lambda$, is independent of the cell detachment rate $k_d$ while it has a non-trivial dependence on the allocation towards chitinase synthesis $\varphi_E$.

To further understand this behavior, let's first consider the simplifying case in which $k'_a = 0$. This more closely resembles the situation that these types of system may encounter in the ocean since any detached enzymes would easily diffuse away. In this case, Eq. S30 becomes:

$$\lambda(\varphi_E) = (k_E Y_b \varphi_E - k'_d)/2 + |\, k'_d - \kappa_E Y_b \varphi_E| \\ = \begin{cases} 0 & \text{for } k'_d > k_E Y_b \varphi_E \\ k_E Y_b \varphi_E - k'_d & \text{for } k'_d < k_E Y_b \varphi_E \end{cases} \quad \text{(S31)}$$

There is a minimum value of $\varphi_E$ below which there can be no growth: $\varphi_{E,min} = k'_d/(\kappa_E\, Y_b)$. Note that if $k'_d = 0$ (i.e: the chitinases never detach) then $\lambda > 0$ and there is no threshold behavior: the culture can always grow. In simpler terms, if enzymes remain on particles and actively produce nutrients, a growing steady-state can always be obtained.

While the overall growth rate $\lambda(\varphi_E)$ shows no $k_d$ dependence, the replication rate of cells on particles, $r_s$, on the other hand varies with the detachment rate, $k_d$ since $r_s = \lambda + k_d$. Given the form of $r_s$ from Scott et. al (2010) described in Eq. S9, there will be a maximum $r_s$ achieved after which the dominating term will come from chitinase overexpression. To see this, we translate the result for $\lambda$ in Eq. S31 to $r_s$ and the nutrient concentration $n/K$. Given Eq. S9:

$$r_s = r_{max}\frac{n/K}{1+n/K}\left(1 - \frac{\varphi_E}{\varphi_{max}}\right)$$

$$\rightarrow \frac{n}{K} = \frac{\lambda + k_d}{r_{max}}\left((1 - \varphi_E/\varphi_{max}) - \frac{\lambda + k_d}{r_{max}}\right)^{-1} \quad \text{(S32)}$$

Let $\frac{n^*}{K}$ be the value at which Eq. S32 diverges. This corresponds to $\varphi_E^* = \varphi_{max}(1 - (\lambda + k_d)/r\_max)$. Therefore:

$$r_s(\varphi_E, k_d) = \begin{cases} \lambda(\varphi_E) + k_d & \text{for } \varphi_E < \varphi_E^* \\ r_{max}\left(1 - \frac{\varphi_E}{\varphi_{max}}\right) & \text{for } \varphi_E > \varphi_E^* \end{cases} \quad \text{(S33)}$$

Similarly, for $\lambda$ we have:

$$\lambda(\varphi_E, k_d) = \begin{cases} k_E Y_b \varphi_E - k'_d & \text{for } \varphi_E < \varphi_E^* \\ r_{max}\left(1 - \frac{\varphi_E}{\varphi_{max}}\right) - k_d & \text{for } \varphi_E > \varphi_E^* \end{cases} \quad \text{(S34)}$$

In principle, the population could increase its chitinase excretion rate until it reaches $\varphi_E^*$. However, for the range of parameters we experimentally observe, we find that $\varphi_E$ is kept rather small by the cells. Specifically, $\varphi_E^* \approx 15\%$ while the measured value of $\varphi_E \approx 3\%$. We speculate that this may be a mechanism to keep nutrient levels $\frac{n}{K}$ low which would result in a tradeoff between $k_d$ and $\lambda$. For a fixed nutrient concentration $n/K$ keeping $\varphi_E$ constant, we find this tradeoff as:

$$\rightarrow \lambda + k_d = r_{max}\left(1 - \frac{\varphi_E}{\varphi_{max}}\right)\frac{n/K}{1+n/K} \quad \text{(S35)}$$

*c- An easier derivation*

In this section, we focus on the flux balance between nutrient generation and uptake by replicating cells. Our goal is to give a more intuitive derivation for $\lambda(\varphi_E)$ in the regime in which $\varphi_E < \varphi_E^*$.

The fraction of the total nutrient flux allocated to chitinase synthesis in steady-state is $\varphi_E = J_E/J_n$ , where $J_E Y_b = \lambda(\varepsilon_b + \varepsilon_s)$. We rearrange this equation to get $\lambda(\varphi_E)$:

$$\lambda = (\varphi_E J_n Y_b)/(\varepsilon_b + \varepsilon_s) \quad \text{(S36)}$$

Eq. S19 allows to explicitly express the total nutrient flux $J_n$ in terms of the other parameters:

$$\lambda = \varphi_E k_E Y_b\, (\varepsilon_s/(\varepsilon_b + \varepsilon_s\,)) \quad \text{(S37)}$$

The population increase rate $\lambda$ only depends on enzyme amounts and not cellular parameters. In particular, it is independent of the detachment rate $k_d$. Intuitively, this is the simple statement that the rate at which the total population increases only depends on nutrient generation by chitinases regardless of the population partioning between the planktonic and surface asscoiated phase. The ratio $\eta' \equiv \varepsilon_b/\varepsilon_s = k_d'/(\lambda + k_a')$ is determined by Eq. S17, and allows to rewrite $\lambda$ as:

$$\lambda = \varphi_E k_E Y_b(\lambda + k_a')/(\lambda + k_a' + k_d') \quad \text{(S38)}$$

We clearly see that if $k_a' = 0$, we recover the previous result Eq. S34:

$$\lambda + k_d' = \varphi_E k_E Y_b \quad \text{(S39)}$$

In the most general case where $k_a' \neq 0$, we can solve the quadratic equation S38 and recover the general result in Eq. S29.

Given the enzyme dynamics in this broadcast case are decoupled from cellular dynamics, this offers an intuitive reason as to why $\lambda$ is independent of $k_d$. The linear form of the dependence is akin to the ribosomal growth law observed in many organisms. It states that if the main bottleneck in growth is nutrient production, then the extent of chitinase secretion determines the growth rate, since it is responsible for dialing nutrient production.

Moreover, the independence of $\lambda$ from $k_d$ suggests a surprising result: that the system can get away with arbitrarily high detachment rates as long as it's secreting chitinases. This independence of the growth rate from the detachment rate is enabled by the feedback on the surface of the particles between the replication rate of cells and their detachment rate. Fewer cells on particles due to a high detachment rate implies that the nutrient concentration per cell is increased thus resulting in a higher replication rate on the particles. This feedback, facilitated by chitinase secretion and accumulation on the particles stabilizes the system.

*d- <u>Some key takeaways:</u>*

- A minimum chitinase production amount is required to overcome the detachment of chitinases $k'_d$ and for a steady-state to be possible. This threshold for growth is $\varphi_{E,min} = k'_d/(\kappa_E\, Y_b\,)$
- The overall population increase rate λ depends linearly on $\varphi_E$, the allocation towards chitinase production, until $\varphi_E = \varphi_E^*$. At this point, protein overproduction becomes costly, as the nutrient concentration is above what's needed to attain the maximum growth rate $r_{max}$.
- For a fixed value of $\varphi_E$, $\lambda$ is independent of $k_d$ for $\varphi_E < \varphi_E^*$. This is because in the case of broadcast enzymes, the enzymes' dynamics are decoupled from the cells' dynamics. For $\varphi_E > \varphi_E^*$ it decreases linearly with the detachment rate, $k_d$.
- For a fixed value of $k_d$, $\lambda$ increases linearly with $\varphi_E$ until it reaches its peak value at $\varphi_E = \varphi_E^*$. After this point, the cost of protein overproduction causes $\lambda$ to decrease linearly with $\varphi_E$.
- Increasing chitinase production such that $\varphi_E = \varphi_E^*$ comes with the disadvantage of increasing the nutrient concentration on the surface of particles, since $\varphi_E^*$ is defined as the chitinase production level at which the nutrient concentration $n/K$ diverges.

#### 3- <u>Case of cell-bound enzymes</u>

*a- <u>Constraints on the parameters</u>*

To outline the benefits of chitinase excretion uncovered in the previous section, we use our general framework to explore the case in which chitinases remain bound to the cell wall as a contrasting scenario. This means that the enzymes' detachment and attachment rates correspond to those of the cells and that the enzyme production rate is the same as the surface-associated cells' replication rate:

$$k'_a = k_a = 0\,,\; k'_d = k_d \text{ and } \beta_s = r_s^{bound} \qquad \text{(S40)}$$

Given these constraints on the parameters, we rewrite the general population level model in Eqs. S1-S5 including the experimental findings described in Section 2-a as:

$$\frac{d\rho_b}{dt} = k_d\, \rho_s \qquad (S41)$$

$$\frac{d\rho_s}{dt} = (r_s^{bound} - k_d)\rho_s \qquad (S42)$$

$$\frac{d\varepsilon_b}{dt} = k_d\, \varepsilon_s \qquad (S43)$$

$$\frac{d\varepsilon_s}{dt} = r_s^{bound}\rho_s - k_d\, \varepsilon_s \qquad (S44)$$

$$\frac{dn}{dt} = k_E\varepsilon_s - \mu\rho_s \qquad (S45)$$

In this case, we see that the equations for $\varepsilon_i$ are redundant. Their dynamics are related to those of the $\rho_i$ by a simple ratio. Moreover, we can easily solve this system since all of the variables depend on the dynamics of $\rho_s$ only, which can in turn be trivially solved. In steady-state our model reduces to the following set of equations written in terms of $\lambda^{bound}$, the exponential growth rate:

$$\eta \equiv \rho_b/\rho_s = k_d/\lambda^{bound} \qquad (S46)$$

$$\lambda^{bound} = r_s^{bound} - k_d \qquad (S47)$$

$$\eta' \equiv \varepsilon_b/\varepsilon_s = k_d/\lambda^{bound} = \eta \qquad (S48)$$

$$\varepsilon_s/\rho_s = (r_s^{bound})/(\lambda^{bound} + k_d) \qquad (S49)$$

$$k_E\varepsilon_s = \mu\rho_s = J_n \qquad (S50)$$

*b- <u>Examining the effect of $\varphi_E^{bound}$ and $k_d$ on the system</u>*

We first examine the effect of the detachment rate $k_d$ on the overall population increase rate $\lambda^{bound}$ and replication rate $r_s$. Let us denote the allocation towards chitinase synthesis in this case as $\varphi_E^{bound}$.
In steady-state, the flux dedicated to chitinase synthesis $J_E = \varphi_E^{bound} J_n = \varphi_E^{bound} k_E\varepsilon_s$ is:

$$J_E Y_b = \Sigma_i \dot{\varepsilon_i} = r_s^{bound}\rho_s \qquad (S51)$$

This leads to the simple relation between the replication rate $r_s$ and the allocation towards chitinase synthesis:

$$r_s^{bound} = \varphi_E^{bound} k_E Y_b \varepsilon_s/\rho_s \qquad (S52)$$

Substituting this expression for the replication rate in Eq. S47, we get:

$$\lambda^{bound}\left(\varphi_E^{bound}, k_d\right) = k_E Y_b \varphi_E^{bound} - k_d \qquad (S53)$$

And thus

$$r_s^{bound}\left(\varphi_E^{bound}\right) = k_E Y_b \varphi_E^{bound} \qquad (S54)$$

Including the effect of chitinase overproduction through Eq. S9, allows to find:

$$r_s^{bound} = r_{max}\left(\frac{n/K}{1+n/K}\right)\left(1-\varphi_E^{bound}/\varphi_{max}\right)$$

Solving for the nutrient concentration $n/K$ , we find:

$$\rightarrow n/K \ = \left(\frac{r_{max}}{r_s}\left(1-\varphi_E^{bound}/\varphi_{max}\ \right)-1\right)^{-1} \quad \text{(S55)}$$

Substituting for the value of the replication rate in Eq. S52, the nutrient concentration in this expression diverges when:

$$\varphi_E^{*bound} = \left(k_E\ Y_b/r_{max} + 1/\varphi_{max}\ \right)^{-1}$$

The full form solution for the replication rate $r_s$ is thus:

$$r_s^{bound}\left(\varphi_E^{bound}, k_d\right) = \begin{cases} \varphi_E^{bound} k_E Y_b & \text{for } \varphi_E^{bound} < \varphi_E^{*bound} \\ r_{max}\left(1-\frac{\varphi_E^{bound}}{\varphi_{max}}\right) & \text{for } \varphi_E^{bound} > \varphi_E^{*bound} \end{cases} \quad \text{(S56)}$$

The overall population replication rate is:

$$\lambda^{bound}(\phi_E, k_d) = \begin{cases} \varphi_E^{bound} k_E Y_b - k_d & \text{for } \varphi_E^{bound} < \varphi_E^{*bound} \\ r_{max}\left(1-\frac{\varphi_E^{bound}}{\varphi_{max}}\right) - k_d & \text{for } \varphi_E^{bound} > \varphi_E^{*bound} \end{cases} \quad \text{(S57)}$$

*c- Comparison between the broadcast and cell-attached case*

- In both cases, there is a threshold value of chitinase expression below which no exponential growth can occur. In the case of secreted enzymes, this threshold depends on the chitinases' detachment rate $k_d'$, while in the case of bound enzymes it depends on the cell detachment rate $k_d$.
- The experimental observation that $k_d/k_d' \approx 10$ illustrates the added difficulty of achieving exponential growth when enzymes remain cell-bound.
- We note that for the same level of chitinase expression $\varphi_E^{bound} = \varphi_E$, the exponential growth rate in the case of secreted enzymes is always higher, such that: $\lambda \geq \lambda_s^{bound}$. The two rates are equivalent if there is no detachment rate.
- Overall, enzyme secretion allows for higher detachment rates of the cells without affecting the overall growth rate.

#### 4- A conflict between privatization and high detachment

Our experimental observations demonstrate that the chitin expression level is kept low $\varphi_E \approx 3\%$. A possible rationale against increasing this level further is that it would lead to higher concentrations of labile nutrients (GlcNAc) on the particle surface (Fig. 4d), which could in turn promote the growth of cheaters since chitin particles are complex ecological systems[6–9]. Thus, maximizing the population increase rate may not advantage 1A01's long-term survival strategy.

Why don't cells thus "privatize" their resources as a way of bypassing cheaters who may feast on these "public goods"[10,11] by binding them to the cell surface[12–15] such that nutrients generated are immediately taken up? The model outlined above in section 3 examines this scenario (Fig. N-I).

In this case, the observed chitinase expression for 1A01 ($\varphi_E \approx 3\%$) would give rise to a replication rate of $r_s^{bound} \approx 0.06\, h^{-1}$, well below the detachment rate observed $k_d \approx 0.2\, h^{-1}$ , thus failing to result in exponential growth. Generally, the overall population increase rate in the case of bound-enzymes (Fig. N-I b) is always lower than in the case of broadcast enzymes (Fig. N-I c).

The physiological reason cells with bound enzymes would experience slower rates in their population increase is that when they detach from particles, these cells carry their bound enzymes with them (Fig. 5a). Since chitinases are only useful when in contact with their substrate, this is a wasteful strategy. In contrast, when enzymes are secreted and released, they remain on particles even as cells detach[16]. This illustrates a basic conflict between binding enzymes to cells and achieving high detachment rates.

To reach a comparable growth rate as 1A01 (red dotted line in Fig. N-I b), cells with bound chitinases would need to increase the allocation for chitinase expression 4-fold, to $\varphi_E \approx 12\%$ of the nutrient influx. This corresponds to ~20% of the cellular proteome on particles (Extended Data Table 1), making it comparable to the entire ribosome content of the cell (Fig. 4f). Such high chitinase expression would deprive cells from expressing a variety of other proteins, e.g., motility, Type-VI secretion system, etc. (Table S1), which can enable the adoption of alternative survival strategies[17].

#### 5- On the attachment rate

While our experiments suggest that in the steady-state, the attachment rate $k_a \ll \lambda$, the initial dynamics of the system must rely on an attachment rate that's high enough to get the culture started. Our experiment in Fig.1d where planktonic cells are re-incubated in a fresh culture with fresh chitin particles is surprising in this sense, since we observe no lag time. Is this lack of lag time consistent with our estimate for the attachment rate?

We first provide an order of magnitude estimate for the lag based on the attachment rate. Assuming encounters between cells and particles are diffusion-limited, then the rate $\Gamma$ for a cell to encounter a particle of radius $R$ is given by

$$\Gamma = 4\pi D R \rho_c$$

where $D$ is the effective diffusion coefficient of randomly tumbling 1A01 cells and $\rho_c$ is the particle density. For our chitin samples, $R \approx 150\, \mu m$ and $\rho_c \approx 3000/mL$. Also, $D \approx 100\, \mu m^2/s$ based on our measurement of 1A01's swimming characteristics. These numbers lead to an encounter rate of $\Gamma \approx 2/h$. The attachment rate $k_a$ is given by

$$k_a = s \cdot \Gamma$$

where $s$ is the "stickiness", i.e., the probability that a cell attaches to the particle after encounter. The value of this parameter in the literature range between $1 - 10\%$ [18,19]. Assuming the stickiness $s$ for 1A01 is on the low end, i.e., $s = 1\%$, then $k_a \approx 0.02/h$ which is barely consistent with the condition $k_a \ll \lambda$.

Next, we note that during the initial stage after inoculation, the dominant process is attachment. If we discard the effect of both growth and detachment, the surface-associated density is:

$$\rho_s(t) = \rho_b(0)(1 - e^{-k_a t})$$

For 10% of the initial population to adhere to the particles given the above parameters, it would take:

$$t^* \approx 5h$$

which is faster than the 12h timescale for the growth of our culture and therefore not easily noticeable in the data of Fig. 1b.

However, if it turns out that $s \ll 1\%$, then $t^* \gg 5h$, and a lag should be readily noticeable.

Experimentally, we can think of a number of practical effects making it difficult to observe a significant lag upon re-incubation of the planktonic component of the chitin culture into fresh chitin media (Fig. 1b):

1- Planktonic cells produce some amount of chitinases which help get the culture started since we observe a small amount of chitinases in their proteome (Table S1).
2- Some small particles containing growing particle-associated cells (which were impossible to exclude completely) were transferred to the fresh chitin culture and they presented an immediate source of growth.

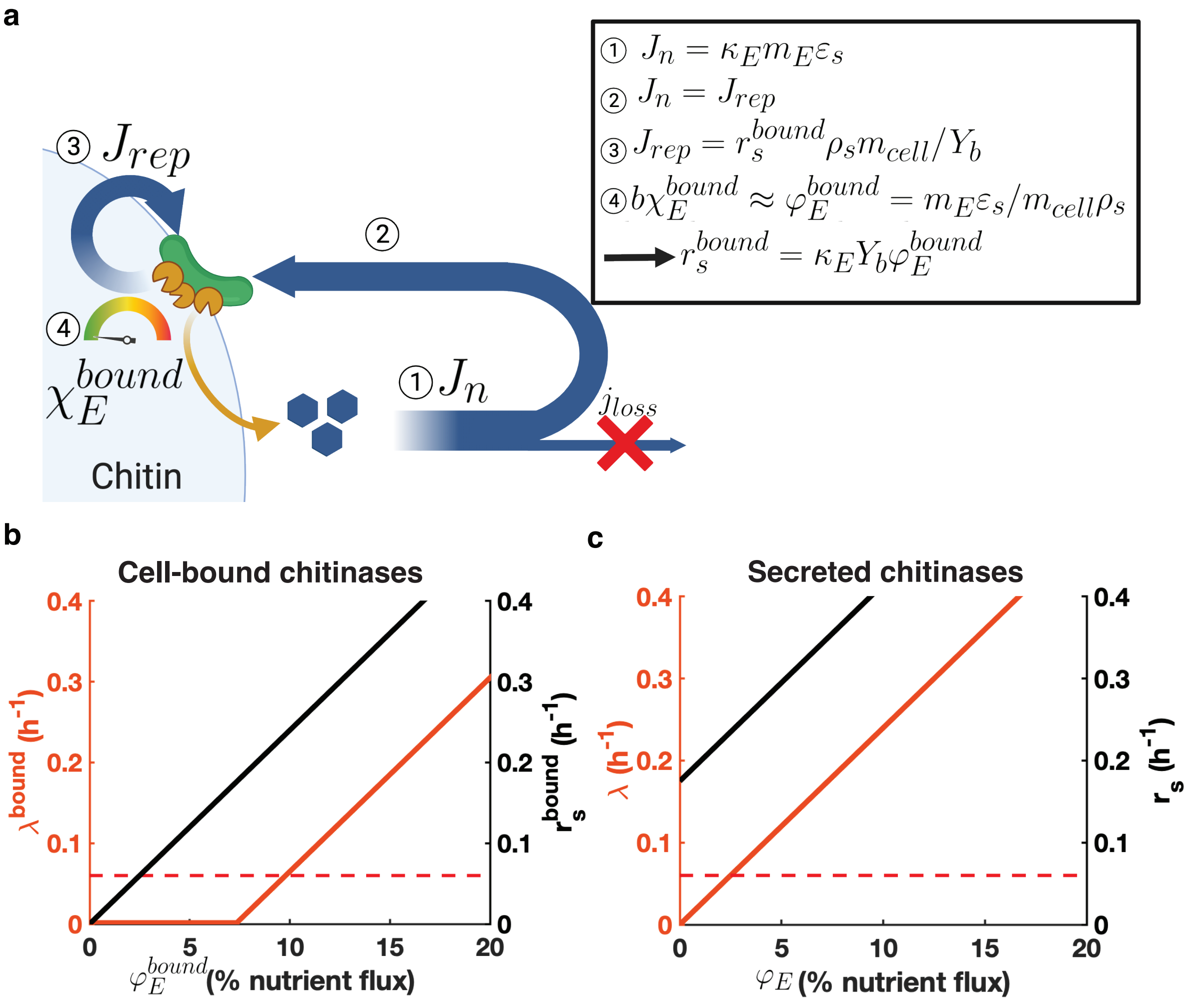


**Figure N-I: The case of enzymes bound to the cell surface. a)** Model describing chitinase synthesis and labile nutrient generation by chitinases which are bound to the cell surface. ① Enzymes bound to the surface of the particles, of concentration $\varepsilon_s$ (yellow pacmans), produce GlcNAc molecules (blue hexagons) at a rate $m_E \kappa_E \varepsilon_s$, where $\kappa_E$ is the catalytic rate of the chitinases per enzyme mass and $m_E$ the enzyme mass. ② The total flux of generated nutrients $J_n$ fully goes towards cellular biomass production $J_{rep}$ since the loss of nutrients due to diffusion $j_{loss}$ is negligible during steady state growth; see text. ③ The nutrient flux related to growth is proportional to the replication rate of the cells such that $J_{rep} = r_s^{bound} \rho_s m_{cell}/Y_b$ where $r_s^{bound}$ is the replication rate of surface associated cells, $\rho_s$ their density and the factor $m_{cell}/Y_b$ simply a conversion factor between biomass and nutrient concentration. ④ The fraction of the total flux $J_n$ allocated towards chitinase synthesis is $\chi_E^{bound}$ and can be dialed by the cells. It can be expressed as $b\chi_E^{bound} = \varphi_E^{bound} \approx m_E \varepsilon_s / m_{cell} \rho_s$ where $b = m_p/m_{cell}$.Taken together, relations ① through ④ lead to the equation describing the replication rate of surface associated cells Eq. (8) $r_s^{bound} = \kappa_E Y_b \varphi_E^{bound}$. **b)** Growth rate $\lambda^{bound}$ (left, red axis) and replication rate of surface associated cells $r_s^{bound}$ as a function of the chitinase fraction $\chi_E^{bound}$ in the case of cell-bound enzymes for the measured detachment rate, $k_d = 0.18h^{-1}$. In this case, it is the replication rate that is proportional to the chitinase fraction whereas the overall population increase rate is reduced by the detachment rate. We see that at the chitinase level measured for 1A01 $\varphi_E = 3\%$, the system cannot achieve an exponentially increasing steady-state. To achieve a similar population increase rate than 1A01, $\lambda^{bound} = 0.06h^{-1}$ (indicated by the dotted red line), cells would need to express ~3 times as many chitinases $\varphi_E^{bound} \approx 10\%$. **c)** As a comparison, we once again plot the growth rate (left, red axis) and replication rate (right black axis) for the case of secreted enzymes as a function of chitinase expression for the measured detachment rate $k_d = 0.18h^{-1}$ . The red dotted line corresponds to the observed growth rate $\lambda = 0.06h^{-1}$. Comparing Panels b and c, we see that for a similar chitinase expression level where $\varphi_E^{bound} = \varphi_E$, $\lambda^{bound}$ is always lower than $\lambda$.

#### II- On chitinase dynamics

In this section we explain how the parameters governing chitinase dynamics (Figs. 4 and Extended Data Fig.6 in the main text) were determined. Experiments consisted of extracting samples from a steady-state growing culture at different planktonic ODs. These samples were then treated with chloroform to disable nutrient uptake by the cells, which allowed to directly monitor the production of nutrients. Control experiments showed that chloroform indeed disabled cell growth and GlcNAc uptake but that it had a detrimental effect on the activity of purified enzymes. This effect is taken into account for assessing the catalytic rate of the enzymes (see Methods).

We measured GlcNAc concentration accumulation traces over time given an initial planktonic OD. To interpret this data, we used a simple model in which two pools of enzymes, planktonic enzymes $\varepsilon_b$ and enzymes attached to the surface of the particles $\varepsilon_s$ can attach and detach with rates $k'_a$ and $k'_d$ . Only enzymes attached to the surface of the particles can produce nutrients with a catalytic rate $k_E$.

This corresponds to Eqs. S1-S5 in the case where $\rho_b = \rho_s = 0$:

$$\frac{d\varepsilon_b}{dt} = k'_d\varepsilon_s - k'_a\varepsilon_b \quad \text{(S58)}$$

$$\frac{d\varepsilon_s}{dt} = k'_a\varepsilon_b - k'_d\varepsilon_s \quad \text{(S59)}$$

$$\frac{dn}{dt} = k_E\varepsilon_s \quad \text{(S60)}$$

Our goal is to solve for the full dynamics of $n(t)$. Let $\varepsilon_{tot} = \varepsilon_b + \varepsilon_s$ be the total amount of chitinases in the system:

$$\frac{d\varepsilon_{tot}}{dt} = \frac{d\varepsilon_b}{dt} + \frac{\varepsilon_s}{dt} = 0 \quad \text{(S61)}$$

The conservation of $\varepsilon_{tot}$ allows to reduce the system in terms of one variable only:

$$\frac{d\varepsilon_s}{dt} = k'_d(\varepsilon_{tot} - \varepsilon_s) - k'_d\varepsilon_s \quad \text{(S62)}$$

$$\frac{dn}{dt} = k_E\varepsilon_s \quad \text{(S63)}$$

Eq. S62 can be easily solved given the initial condition $\varepsilon_s(0)$:

$$\varepsilon_s(t) = \frac{k_a{}'\varepsilon_{tot}}{k_a{}' + k_d{}'} + \left(\varepsilon_s(0) - \frac{k_a{}'\varepsilon_{tot}}{k_a{}' + k_d{}'}\right) e^{-(k'_a + k'_d)t} \quad \text{(S64)}$$

We then integrate Eq. S64 to find the solution of Eq. S63 given the initial condition $n(0) = 0$ to find:

$$\frac{n(t)}{k_E} = \frac{k_a{}'\varepsilon_{tot}}{k_a{}' + k_d{}'} t - \left(\frac{\varepsilon_s(0)}{k_a{}' + k_d{}'} - \frac{k_a{}'\varepsilon_{tot}}{(k_a{}' + k_d{}')^2}\right)\left(e^{-(k'_a + k'_d)t} - 1\right) \quad \text{(S65)}$$

Let us consider various limits of Eq. S65. At short times, (i.e: $t \ll \frac{1}{k_a{}' + k_d{}'}$):

$$\lim_{t\to 0} \frac{n(t)}{k_E} = \frac{k_a{}'\varepsilon_{tot}}{k_a{}'+k_d{}'} t + \left(\frac{\varepsilon_s(0)}{k_a{}'+k_d{}'} - \frac{k_a{}'\varepsilon_{tot}}{(k_a{}'+k_d{}')^2}\right)(k_a' + k_d')t$$
$$= \varepsilon_s(0)t \qquad \text{(S66)}$$

Initially, the GlcNAc concentration increases linearly with a slope $s_1 = k_E \varepsilon_s(0)$, allowing to estimate the value of $k_E$.

At longer times, (i.e: $t \gg \frac{1}{k_a{}'+k_d{}'}$ ):

$$\lim_{t\to\infty} \frac{n(t)}{k_E} = \frac{k_a{}'\varepsilon_{tot}}{k_a{}'+k_d{}'} t + \left(\frac{\varepsilon_s(0)}{k_a'+k_d'} - \frac{k_a'\varepsilon_{tot}}{(k_a'+k_d')^2}\right) \qquad \text{(S67)}$$

The GlcNAc concentration increases linearly with a slope $s_2 = k_E \frac{k_a'\varepsilon_{tot}}{k_a'+k_d'}$. Taking the ratio of the two slopes allows us to determine the ratio $k_d'/k_a'$ since:

$$\frac{s_1}{s_2} = \varepsilon_s\,(0)/\varepsilon_{tot}(1 + k_d'/k_a') \qquad \text{(S68)}$$

The time scale at which $n(t)$ changes from one slope to the other allows to fix $k_a' + k_d'$ and determine all parameters. We notice that the two lines described in Eqs. S66 and S67 intersect exactly at $t^* = 1/(k_a' + k_d')$

$$\varepsilon_s(0)t^* = \frac{k_a'\varepsilon_{tot}}{k_a' + k_d'} t^* + \left(\frac{\varepsilon_s(0)}{k_a' + k_d'} - \frac{k_a'\varepsilon_{tot}}{(k_a' + k_d')^2}\right)$$
$$\to t^* = (k_a' + k_d')^{-1} \qquad \text{(S69)}$$

Given the estimates for $\varepsilon_s(0)$ and $\varepsilon_{tot}$, we determine the value of the enzyme parameters $k_a'$, $k_d'$ and $k_E$.

#### III- Spatial model for growth on a chitin particle

A key assumption made in our population-level model (Supp. I) is that the flux of nutrients is balanced: all nutrients generated are used for either biomass or chitinase production and none are lost to the surrounding environment. This assumption which was backed by our data (Fig. 2, ED Figs. 2,3) allowed us to collapse the spatial structure into a simple two-compartment model: planktonic vs. particle-associated cells and enzymes.

Theoretically, this assumption is justified because nutrient generation, which is proportional to the chitinase concentration on particles, and the nutrient uptake, which is proportional to the number of cells on particles, are both increasing exponentially, whereas the nutrient leakage is finite (because the nutrient concentration on the surface does not increase with time). Therefore, in the long-time limit, nutrient leakage will be an exponentially small fraction of the flux of nutrient generation and uptake.
Below, we provide a simplified spatial model to see how this balance quantitatively works out. In the more realistic case, cell colonies on the particles may be patchy (ED Fig.4) and these patches may not be co-localized with the enzymes. However, such mismatches are unlikely to last long since cells would chemotax to the vicinity of nutrients [20]and they will then replicate more rapidly until the local nutrient generation and uptake fluxes are balanced.

##### 1- Case with a constant planktonic cell density

Let us consider a spherical chitin particle of radius $R_0$ coated with a monolayer of cells at the surface with density $\sigma$. A similar model is developed in Nguyen et al. (2021)[21]. GlcNAc is generated at the surface with a rate $k_E \varepsilon_s$ where $k_E$ is the catalytic rate of the enzymes as determined above and $\varepsilon_s$ is the active enzyme concentration on the surface of the particles. Away from the particles, let's consider planktonic cells with uniform density $\rho_b$.

Both surface associated and planktonic cells uptake nutrients following Monod kinetics with growth rate $r(n(R)) = \frac{r_{max} n(R)}{K_n Y}$ where $R$ is their radial position away from the center of the particle and $Y$ is the yield of 1A01 cells growing on GlcNAc monomers.

From this formulation, it is clear that the nutrient gradient determines the replication rates of the two cellular subpopulations. The nutrient concentration in the system is:

$$\frac{\partial n(R,t)}{\partial t} = D_n \nabla^2 n(R,t) + k_E \varepsilon_s(t)\delta(R-R_0) - \frac{r_{max} n(R_0,t)}{K_n Y}\sigma(t)\delta(R-R_0) - \frac{r_{max} n(R,t)}{KY}\rho_b(\mathrm{t})\Theta(R-R_0) \quad \text{(S70)}$$

where $\delta$ is the Dirac delta function and $\Theta$ is the Heaviside step function.

By assuming instantaneous equilibration of the nutrient field (i.e: $\frac{\partial n(R,t)}{\partial t} = 0$), writing the Laplacian in spherical coordinates as $\nabla^2 n(R) = \frac{1}{R}\frac{d^2}{dR^2}(R \cdot n(R))$ and defining $g(R) = R \cdot n(R)$, we solve the following equation for the nutrient field:

$$D_n g''(R) + R k_E \varepsilon_s \delta(R - R_0) - \frac{r_{max} g(R_0)}{K_n Y} \sigma \delta(R - R_0) - \frac{r_{max} g(R)}{K_n Y} \rho_b \Theta(R - R_0) = 0 \quad \text{(S71)}$$

For the region $R > R_0$, this simplifies to:

$$g''(R) = \frac{r_{max} \rho_b}{D_n K_n Y} g(R) = \kappa^2 g(R) \quad \text{(S72)}$$

Where we define $\kappa^2 = \frac{r_{max} \rho_b}{D_n K_n Y}$ such that $\kappa^{-1}$ is the screening length.

The boundary condition at $R_0$ is found by integrating Eq. S71 in an infinitesimal region around $R_0$:

$$D_n (g'(R_0^+) - g'(R_0^-)) + R_0 k_E \varepsilon_s - \frac{r_{max} g(R_0)}{K_n Y} \sigma - \frac{r_{max}}{K_n Y} \rho_b (g(R_0^+) - g(R_0^-)) = 0 \quad \text{(S73)}$$

To prevent inward diffusive flux we have that $g'(R_0^-) = n_0$, and $g(R_0^+) = g(R_0^-)$ to insure continuity. Simplifying the above equation, we get:

$$D_n g'(R_0) - D_n n_0 + R_0 k_E \varepsilon_s - \frac{r_{max} g(R_0)}{K_n Y} \sigma = 0 \quad \text{(S74)}$$

$$g'(R_0) - n_0 + \frac{R_0}{D_n} k_E \varepsilon_s - \frac{r_{max} g(R_0)}{D_n K_n Y} \sigma = 0 \quad \text{(S75)}$$

The solution to Eq. S74 where $g(R \to \infty)$ is bounded is:

$$g(R) = g_0 e^{-\kappa(R - R_0)} \quad \text{(S76)}$$

In terms of the nutrient concentration explicitly:

$$n(R) = \frac{n_0 R_0}{R} e^{-\kappa(R - R_0)} \quad \text{(S77)}$$

Where $n_0$ is obtained by solving the boundary condition

$$n_0 = \frac{k_E \varepsilon_s}{D_n (\kappa + R_0^{-1}) + \frac{r_{max}}{K_n Y} \sigma} \quad \text{(S78)}$$

Note that for $\kappa^{-1} \gg R_0$ we have $n_0 = \frac{R_0 k_E \varepsilon_s}{D_n (1 + \frac{r_{max}}{K_n Y} \sigma)}$ and for $\kappa^{-1} \ll R_0, n_0 = \frac{k_E \varepsilon_s}{\kappa D_n}$

In our experiment, $R_0 \approx 150 \mu m$ and the chitin weight density, $\phi_w = 0.2\% w/v$. Given the density of chitin $d_c = 1.4\ g/cm^3$, we estimate the volume density of chitin particles as

$\phi_v = \phi_w/d_c = 1.4 \times 10^{-3}$. This allows to infer the inter-particle spacing $\ell_c = R_0/\phi_v^{1/3} \approx 9R_0$. Moreover, the cell surface density is $\sigma = \left(\frac{\eta R_0}{3\phi_v}\right)\rho_b$ where $\eta = \rho_b/\rho_s$.

The parameters in Table N-III allow us to estimate the screening length: $29\mu m < \kappa^{-1} < 94\mu m$. This is of the order of the particle size, but 10 times smaller than the inter-particle spacing, $l$. We also estimate the value of $n_0$, the nutrient concentration on the surface of chitin particles as $5\ \mu M < n_0 < 20\ \mu M$. This strongly suggests that beyond the small screening distance determined by $\kappa^{-1}$, there is no planktonic replication in the bulk as the nutrient concentration $n(R)$ would quickly fall below the Monod constant, $K_n$, which we take to be $\sim 1\ \mu M$.

| **Parameter** | **Symbol** | **Estimated value** |
|---|---|---|
| Maximum growth rate | $r_{max}$ | $0.8h^{-1}$ |
| Yield on GlcNAc | $Y^{-1}$ | $6.1\ mM/OD_{600}^{plank}$ |
| Monod constant for GlcNAc | $K_n$ | $1\ \mu M$ |
| Diffusion coefficient of GlcNAc | $D_n$ | $600\ \mu m^2/s$ |
| Diffusion coefficient of cells | $D_\rho$ | $100\ \mu m^2/s$ |
| Planktonic density in the culture | $\rho_b$ | $0.05 - 0.5\ OD_{600}^{plank}$ |
| Interparticle spacing | $\ell_c$ | $1.5\ mm$ |

**Table N-III:** Parameters for spatial model of chitin degradation

#### 2- <u>Case with a spatially-dependent planktonic cell density</u>

An important approximation made in the model described above was the constancy of the planktonic cell density. Let us now relax that assumption and allow this cellular field to have spatial structure in response to the spatial nutrient profile. We will analyze the effect of the spatial nutrient profile on the spatial cell density profile and thus reexamine the validity of the constant cell density approximation made above.

Let $\rho(R, t)$ be the planktonic cell density profile. The random tumbling motion of planktonic cells can be effectively described by diffusion at the population level, with a diffusion constant $D_\rho$, while chemotaxis towards nutrients (which can also be chemoattractant[20]) can be described by a concentration-dependent convection term to be detailed below. Additionally, those cells close to the particle can replicate, at a rate that depends on the local nutrient concentration through a Monod form and a Monod constant $K_n$. Similarly to the model above, let $n(R, t)$ be the nutrient profile which diffuses with a constant $D_n$ and is consumed by both planktonic and surface-associated cells. Again, nutrients are produced on the surface of the particles with a rate $k_E \varepsilon_s(t)$. Our system is now comprised of this set of two coupled differential equations:

$$\frac{\partial n(R,t)}{\partial t} = D_n \nabla^2 n(R,t) + \left( k_E \varepsilon_s(t) - \frac{r_{max} n(R_0,t)}{K_n Y} \sigma(t) \right) \delta(R - R_0) - \frac{r_{max} n(R,t)}{K_n Y} \rho(R,t) \Theta(R - R_0) \quad \text{(S80)}$$

$$\frac{\partial \rho(R,t)}{\partial t} = D_\rho \nabla^2 \rho(R,t) + r_{max} \frac{n(R,t)}{n(R,t) + K_n} \rho(R,t) - \chi \nabla \left( \frac{\nabla n(R,t)}{n(R,t) + K_\chi} \rho(R,t) \right) \quad \text{(S81)}$$

We note that in the last term on the right-hand side of Eq. S81, we describe chemotaxis by a modified log-sensing form (or Weber's law, $\nabla n(R,t)/n(R,t)$), with a concentration cutoff $K_\chi$ and with $\chi$ being the chemotactic coefficient. The role of this cutoff is to capture the limited sensitivity of cells to a very low concentration of chemoattractant [22–25]. It plays a crucial role here since in the singular-limit $K_\chi \to 0$, chemotaxis would guide cells from infinitely far away to the particle, even if the nutrient profile attenuates exponentially away from the particle.

Assuming the nutrient field equilibrates instantaneously, i.e., $\frac{\partial}{\partial t} n(R,t) = 0$ (since this timescale is much faster than changes in the planktonic cell density which increase over the time scale $\lambda^{-1}$), Eq. S80 for $R > R_0$ becomes

$$\nabla^2 n(R,t) = \frac{r_{max}}{D_n K_n Y} \rho(R,t) n(R,t), \quad \text{(S82)}$$

which is a nonlinear equation with $\rho(R,t)$ obtained from the solution to Eq. S81, another nonlinear equation. To progress further, we assume that the spatial planktonic cell density profile is only weakly dependent on $R$ and can be considered constant within the millimeter length scale of inter-particle spacing. In other words, we assume $\rho(R,t) \approx \rho(R_0,t)$ as will be self-consistently justified below. With this additional assumption, Eq.(S82) can be solved as described above (Eq. (S71)-(S76)), with the solution

$$n(R,t) = n(R_0,t) \frac{R_0}{R} e^{-\kappa_n(t) \cdot (R - R_0)} \propto e^{-\kappa_n(t) \cdot (R - R_0)} \quad \text{(S83)}$$

where $\kappa_n(t) = \sqrt{\frac{r_{max}}{D_n K_n Y} \rho(R_0,t)}$ is the inverse length scale characterizing the spatial nutrient profile at time $t$. Since $\rho(R_0,t)$ increases exponentially with $\rho(R_0,t) = \rho(R_0)\, e^{\lambda t}$, we see that $\kappa_n^{-1}$, the length scale of the nutrient field, gets exponentially smaller, i.e., $\kappa_n^{-1} \sim e^{-\lambda t/2}$. This means that for exponentially growing cells, the nutrient field gets increasingly localized near the surface of the particle. Similarly to above, for the range of cell densities in our experiments, $0.05 < \rho(t) < 0.5$, we find $29\mu m < \kappa_n^{-1} < 94\mu m$. This length scale, which exponentially decreases in time, is at most 6% of the interparticle spacing ($\ell_c = 1.5mm$).

Given this form of $\kappa_n$ for $n(R,t)$ in Eq. S83, we can ignore both the uptake and chemotaxis term in Eq. S81 for $R \gg \kappa_n^{-1}$. Eq. S81 then becomes:

$$\lambda\rho(R,t) = D_\rho \nabla^2 \rho(R,t) \qquad \text{(S84)}$$

The solution is

$$\rho(R,t) \propto e^{\kappa_\rho (R-R_0)+\lambda t} \qquad \text{(S85)}$$

where $\kappa_\rho^{-1} = \sqrt{D_\rho/\lambda}$. Using parameter values listed in Table N-III, we find $\kappa_\rho^{-1} \sim 2.5\ mm$, which self-consistently justifies the assumption on the weak $R$-dependence we made above. As this is well above the interparticle spacing in our culture, we conclude that the planktonic cell field is delocalized and can be treated approximately as constant.

Although our analysis suggests that the chemotactic term can be ignored since the nutrient concentration quickly drops below the chemotactic sensitivity $K_\chi$, we note that chemotaxis would affect the macroscopic attachment and detachment rates (the parameters $k_a$ and $k_d$ introduced in the main text Eqs. 1-2). Indeed this term tends to localize cells to the vicinity of the particle and thus affect the repartition of surface-associated and planktonic cells near them, thereby modifying the dependence of $k_a$ and $k_d$ on the microscopic attachment and detachment rates that connect $\rho(R_0,t)$ and the density of cells associated with the surface, $N_s(t)$ via boundary conditions for $\rho(R,t)$.

Another effect of chemotaxis is to increase the chitospheric cell density within the distance $\kappa_n^{-1}$. As these cells experience nutrient concentrations similar to that on the particle surface and exchange rapidly with the particle-associated cells, we can regard them together with the cells on particle as "particle-associated" sub-population of cells, referred to as $\rho_s$ in the main text. These modified definitions of $k_a, k_d$, and $\rho_s$ do not affect the model in the main text which is phenomenological in nature. Detailed solution and analysis of the model defined by Eq. S80, S81, and the accompanying boundary conditions will be presented elsewhere.
